# Integrated single-nuclei and spatial transcriptomic profiling of human sacrococcygeal teratomas reveals heterogeneity in cellular composition and X-chromosome inactivation

**DOI:** 10.1101/2025.07.21.665156

**Authors:** Ernesto J. Rojas, Krinio Giannikou, Benjamin J. Huang, Soo-Jin Cho, Marco A. Cordero, Deion Pena, Lan Vu, Aditya Bagrodia, S. Christopher Derderian, Tippi C. MacKenzie, Diana J. Laird

## Abstract

Sacrococcygeal teratomas (SCTs) are the most common neonatal tumors, yet their cellular origins, clinical stratification, and sex bias–occurring three times more in XX than XY individuals—remain poorly understood. To address these gaps, we examined six postnatal (one male and five female) and two prenatal (both female) SCTs by single nuclei RNA-seq and spatial transcriptomics. We identified five broad cellular lineages in SCTs: stroma, epithelia, endothelia, neuroectoderm, and immune. The transcriptomes and lineage compositions showed significant heterogeneity, which offer a framework for future molecular stratification. SCTs are thought to originate from and be propagated by pluripotent cells, notably however, we did not detect these populations. Among female tumors, a subset of cells exhibited biallelic expression of X-linked genes, consistent with X-inactivation failure or reactivation of the once inactivated X-chromosome. These biallelic cells were enriched for developmental and neuronal programs, whereas cells with single-allelic X-chromosome preferentially expressed immune-related genes. Biallelic X-chromosome activation, which can occur only in female cells, may result in transcriptomic features that favor survival of tumor cells, contributing to the sex bias of SCTs. Our findings reveal a link between X-chromosome inactivation and SCT cell identity, suggesting that X-dosage dysregulation may influence SCT heterogeneity and immune landscape.

## Introduction

Sacrococcygeal teratomas (SCTs) are the most common tumors in perinatal patients, arising at the base of the spine and comprising tissues from all three germ layers ^1^. Clinically, SCTs are stratified by neural tissue maturity (mature vs. immature) ^2^, internal vs. external growth (Altman grading) ^3^, and solid vs. cystic composition^4^. Characteristics such as, more immaturity, internal location, or more solid components are associated with poorer outcomes^5,6^ and prenatal resection is required when rapid growth leads to high risk complications ^7^. Epidemiologically, SCTs occur three times more frequently in XX than XY fetuses ^8^, suggesting X-chromosome dynamics during early development may influence susceptibility to tumor development.

The presence of all three germ layers and the prenatal onset of SCTs suggest a potential origin from a pluripotent cell type ^1^. While *in vivo* pluripotency is typically restricted to the preimplantation embryo before gastrulation, primordial germ cells (PGCs) – the embryonic precursor to sex cells – retain a latent pluripotency program that is extinguished during their differentiation into oogonia or pro-spermatogonia in mid-gestation ^9^. In PGCs with two X-chromosomes, one X-chromosome is initially inactivated during peri-implantation, and then reactivated during differentiation into oogonia ^10^. Given that PGCs migrate along the posterior midline of the embryo and that SCTs localize to the coccyx, ectopic (mismigrated) PGCs that escape apoptosis and re-acquire pluripotency have been proposed as a plausible origin of SCTs ^11^. Supporting this hypothesis, pluripotency and PGC markers such as OCT4 and NANOG have been detected in SCTs, and bulk RNA-sequencing of SCTs showed enrichment of PGC-related genes compared to other pediatric tumors ^12–14^.

Investigating SCT inter-tumor heterogeneity has been difficult given their rarity and diverse composition, which limit the discovery of prognostic markers for patient outcomes and the understanding of factors driving tumor phenotypes. To date, the only study of SCT inter-tumor heterogeneity compared mature and immature SCTs by bulk RNA-sequencing ^14^, which showed that mature SCTs expressed more immune signals than immature SCTs, and that pluripotency and PGC markers were detected in both. However, both this bulk RNA-seq study and previous work ^13^ lacked single-cell resolution, precluding the ability to pinpoint cellular coincidence of these markers.

In this study, we used single-nucleus RNA-sequencing (snRNA-seq) and spatial transcriptomics (SpTr) to address these limitations and generate a comprehensive cellular and spatial atlas of eight SCTs. We identified distinct molecular SCT subtypes based on epithelial content and uncovered proliferative mesenchymal and epithelial populations specifically enriched in aggressive prenatal tumors. Contrary to expectations, we found no evidence of a discrete pluripotent cell population. We discovered that cells with two active X-chromosomes (XaXa-like) preferentially express neurodevelopmental genes, while those with one active X-chromosome (XaXi) express immune-related programs, revealing a novel link between X-chromosome inactivation (XCI) state and tumor cell identity. Our findings offer a more granular understanding of the molecular classification of SCTs and provide new directions for investigating tumor heterogeneity, the immune microenvironment, and XCI dysregulation in SCT biology.

## Results

### Single nuclei RNA-sequencing establishes a sacrococcygeal teratoma cell atlas containing lineages from all three germ layers

SCTs are histologically defined by the presence of cell types derived from the three germ layers. Molecular characterization of this heterogeneity could yield valuable insights into tumor biology, prognostic markers, and potential therapeutic targets. We investigated the composition of SCTs with snRNA-seq using 10X Genomics 3’ Chromium platform. Nuclei isolation offers broad representation of cell types from fresh samples, independent of tissue dissociation techniques ^15,16^. We analyzed six postnatally resected SCT samples from patients whose ages ranged from one day to four months (**Fig. 1a**). Our cohort included three predominantly cystic tumors (SCT01, SCT03, SCT10) and three predominantly solid tumors (SCT02, SCT06, SCT09), chosen to reflect the known association between solid components and poorer clinical outcomes ^5^, although the solid tumors in our study have not been associated with adverse outcomes in these patients. We specifically sampled the most solid regions for analysis. To validate our approach, we compared snRNA-seq to single-cell RNA-seq in one SCT sample (SCT01c and SCT01n, **Supplementary Fig 1a,b**), which confirmed a broader diversity of cell types in snRNA-seq (**Supplementary Fig 1a,b**), supporting its use in all subsequent experiments.

**Fig. 1.**
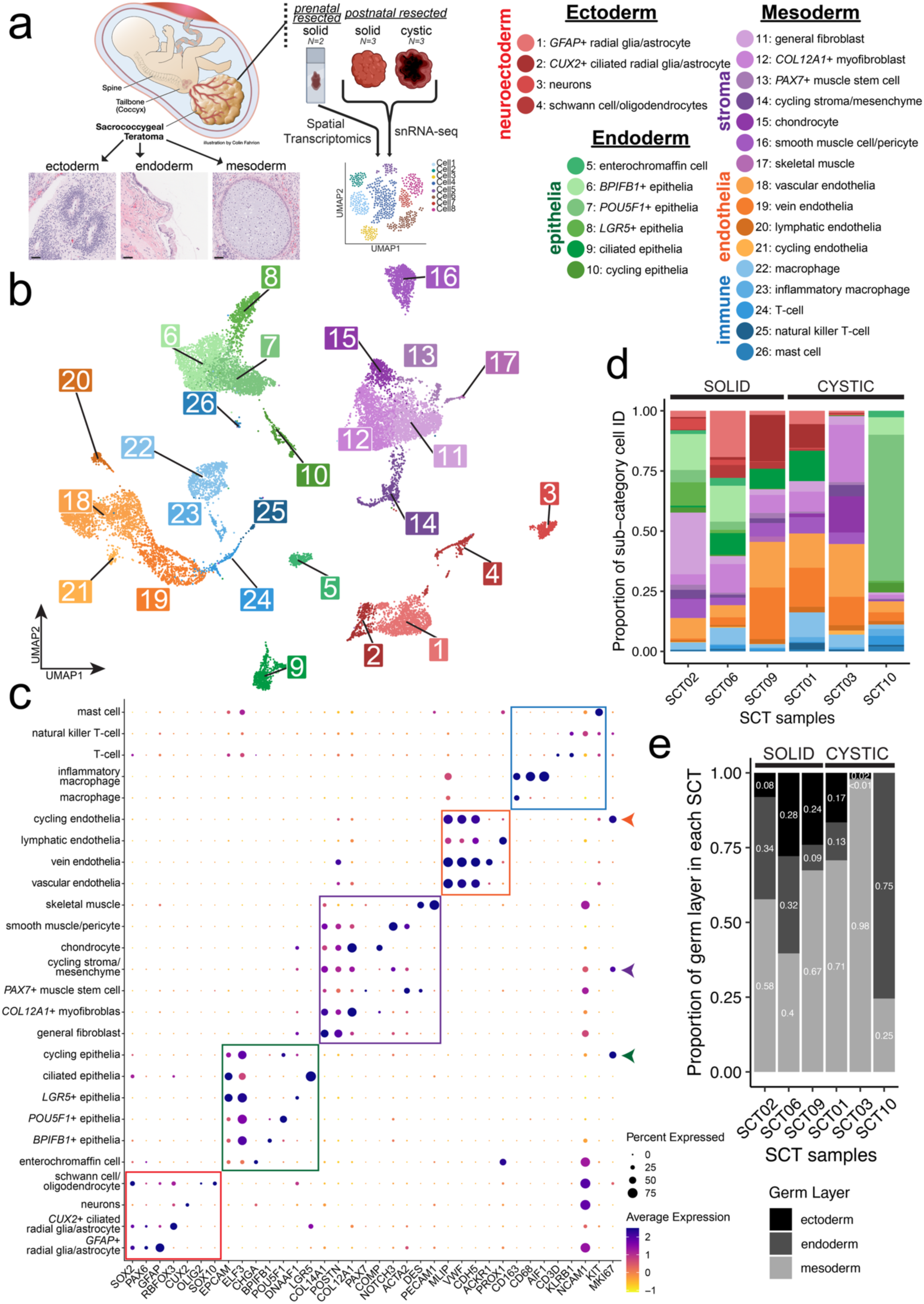
Cell-type composition of an atlas of sacrococcygeal teratomas (SCTs). (a) Experimental design diagram, demonstrating the three germ layers and acquisition of fresh and banked SCTs. Scale bar = 500µm. (b) UMAP projection of *n*=18,034 well-integrated nuclei; cell types are labeled 1-26 and coordinated names are shown in the legend on the right. Colors denote broad categories of cells (red = neuroectoderm, green = epithelia, purple = stroma, orange = endothelia, blue = immune). (c) Dot plot highlighting marker genes for clusters identified; boxes denote markers for each of the broad categories. Cycling cells expressing *MK167* are highlighted with colored arrows on the right side. (d) Bar plot colored by cell identity and grouped by SCT samples; on the left are the 3 predominantly solid tumors; on the right, the 3 predominantly cystic tumors. (e) Bar plot colored by germ layer that the cell type belongs to and grouped by SCT samples.

In addition to postnatal samples, we analyzed two prenatally-resected samples (SCT04, SCT05), both predominantly solid and rapidly growing, which resulted in fetal demise. Because only banked formalin-fixed paraffin-embedded (FFPE) tissue was available, these samples were analyzed by SpTr using 10X Genomics Visium platform. Clinical and pathological characteristics, including patient sex, tumor maturity, and tissue composition are summarized in **Table 1**. Throughout this paper, we refer to tumors resected after birth as SCTs and tumors resected prenatally as prenatally-resected, prSCTs.

**Table 1.** Clinical information of sequenced sacrococcygeal teratomas.

| SCT ID | Sex | Cystic/<br>Solid | Altman<br>Type | Mature/<br>Immature | Size | Age<br>of<br>Birth | Age of<br>Resected | Tissues From<br>Pathology Report |
| --- | --- | --- | --- | --- | --- | --- | --- | --- |
| ***POSTNATAL RESECTION - SINGLE CELL/NUCLEI RNA-SEQ*** |  |  |  |  |  |  |  |  |
| SCT01 | F | cystic | Type 4 | Mature | 5.3 x<br>3.5 x<br>1.5<br>cm | 39w2d | 4 months | neuropil, enteric, upper<br>respiratory epithelia<br>and squamous<br>epithelia |
| SCT02 | F | solid<br>and<br>cystic | Type 4 | Mature | 6.8 x<br>5 x 4<br>cm | 37w1d | 1 month | squamous, enteric,<br>respiratory epithelium<br>and immature<br>mesenchymal<br>components (cartilage,<br>adipose, fibroblast) |
| SCT03 | F | cystic | Type 2 | Mature | 7.9 x<br>7.1 x<br>9.9<br>cm | 36w2d | 1 day | haphazard<br>arrangement of mature<br>tissues including<br>neural, glandular,<br>muscles, adipose<br>tissue, cartilage,<br>respiratory and skin |
| SCT06 | M | mostly<br>solid | Type 3 | Mature | 7.0 x<br>6.2 x<br>4.0<br>cm | 37w | 1 day | heterologous elements,<br>including bone,<br>immature cartilage and<br>adipose,<br>gastrointestinal, glial,<br>adnexal, liver, and<br>respiratory tissue.<br>Mature glial tissue is<br>present at one inked<br>resection margin. No<br>immature neural<br>elements or malignant<br>cells are present. |
| SCT09 | F | solid &<br>cystic | Type<br>2/3 | Mature | 7.5 x<br>6.0 x<br>5.3<br>cm | 38w | 1 day | GFAP+ neural tissue. |
| SCT10 | F | mostly<br>cystic | Type 2 | Mature | 8.5 x<br>4.6 x<br>1.7<br>cm | 38w3d | 3 days | MB1+ areas. Glandular<br>epithelium with<br>inflammatory cells |
| ***PRENATAL RESECTION - SPATIAL TRANSCRIPTOMICS*** |  |  |  |  |  |  |  |  |
| SCT04 | F | solid | Type 3 | Immature | 11 cm | N/A | 20 weeks<br>4 days | immature neuroglial<br>and neuro-epithelial<br>components |
| SCT05 | F | solid | Type 2 | Immature | 6.5 x<br>5.6 x<br>4.2<br>cm | N/A | 15 weeks<br>5 days | a preponderance of<br>immature neural tissue |

Teratomas exhibit diverse differentiation states. To identify and compare cell types across the SCT samples, we used data from human fetal development, stem cell differentiation, and adult teratomas. Cell identities were first annotated independently for each sample and then integrated across tumors. We used four orthogonal approaches to ensure robust classification: (1) automated cell annotation via a hierarchically-organized marker gene map ^17^, (2) correlation of each cluster with published ovarian and human embryonic stem cell (ESC) derived teratoma datasets ^18,19^, (3) projection onto a human fetal development atlas ^20^, and (4) a large-language-model-based classifier ^21^. The four approaches identified overlapping cell types with high concordance (**Supplementary Table 1**). We subsequently integrated the annotations and visualized the results using Uniform Manifold Approximation and Projection (UMAP), demonstrating concordance of cell type annotations across independent samples (**Supplementary Fig 1d**).

We used the cell atlas UMAP to refine the names of the clusters of cells within the tumors. First, we identified five broad cell types of cells types— neuroectodermal, epithelial, stromal, endothelial, and immune (**Fig. 1b**)—which were further resolved into 26 transcriptionally distinct sub-categories using unsupervised clustering paired with canonical and lineage-specific marker genes (**Fig. 1c,d**, **Supplementary Table 2**). The resulting cell identities were consistent with those reported in non-SCT human teratomas (**Supplementary Fig. 1e,f**). Within the neuroectodermal lineage, we identified two astrocyte/radial glia-like populations (*SOX2+, PAX6*+) ^22^: one characterized by high *GFAP+* expression and another *CUX2+* and ciliary marker ^23^, *DNAAF1+*. Other neuroectoderm-derived subtypes included neurons (*RBFOX3+*, protein name NeuN) ^24^ and Schwann cells/oligodendrocytes (*SOX10+*, *OLIG2+*) ^25^. The epithelial lineage was uniformly *EPCAM+* and *ELF3+* ^26^, but further stratified into six transcriptionally distinct populations: enterochromaffin cells identified by the canonical marker *^27^*, *CHGA+*; three populations defined by unique marker genes: *BPIFB1*+, *POU5F1*+ (protein name: OCT4; a canonical marker of pluripotency and germ cell identity), and *LGR5*+ (canonical marker of stem cells in normal colorectal epithelium) ^28^; a fifth ciliary gene, *DNAAF1+* population; and a sixth cycling epithelial population (*MKI67+*). Among the stromal cells, several clusters were classified using canonical markers, including smooth muscle/pericyte-like cells (*ACTA2+*, *NOTCH3+*) ^29^, chondrocytes (*COMP+*) ^30^, skeletal muscle (*DES+, MLIP+*) ^31^, and *PAX7*+ muscle stem cells ^32^. We also identified a population of cycling stromal mesenchyme, a *COL12A1*+ myofibroblast-like population, and a general fibroblast population. Endothelial cells were classified into general vascular (*PECAM1+*, *VWF+*, *CDH5+*), venous (*ACKR1*+), lymphatic (*PROX1+*), and cycling populations ^33^. The immune lineage comprised mast cells (*KIT+*) ^34^, T-cells (*CD3D+*) ^35^, natural killer T-cells (*NCAM1+, KLRB1+*) ^36^, macrophages (*CD68+, CD163+*) ^37,38^ and inflammatory macrophages (*AIFF+*) ^39^. Our results demonstrate that these SCTs comprise cell types from the three primary germ layers (**Fig. 1e**), consistent with established histological analyses of this tumor ^40^. Overall, our SCT cell atlas provides a foundational framework for dissecting tumor heterogeneity in SCTs, highlighting previously unrecognized subpopulations.

### Sacrococcygeal teratomas stratify by transcriptomic profiles driven by cell type composition

A biologic hallmark of SCTs is their extensive cellular heterogeneity. Although most tumors contained cell types from all major germ layer lineages, there were considerable differences across individual tumors (**Fig. 1d,e**). To assess global transcriptional similarities, we applied a pseudo-bulking approach, aggregating scRNA-seq data by tumor and projecting samples into principal component (PC) space (**Fig. 2a**). This analysis revealed three distinct transcriptomic clusters: 1) SCT02 and SCT06; 2) SCT01, SCT03, and SCT09; and 3) SCT10. Spearman correlation analysis further confirmed this grouping, with SCT10 showing the lowest correlation (<0.7) to all other samples, while SCT01, SCT03, and SCT09 formed a highly correlated cluster (>0.8) (**Fig. 2b**).

**Fig. 2.**
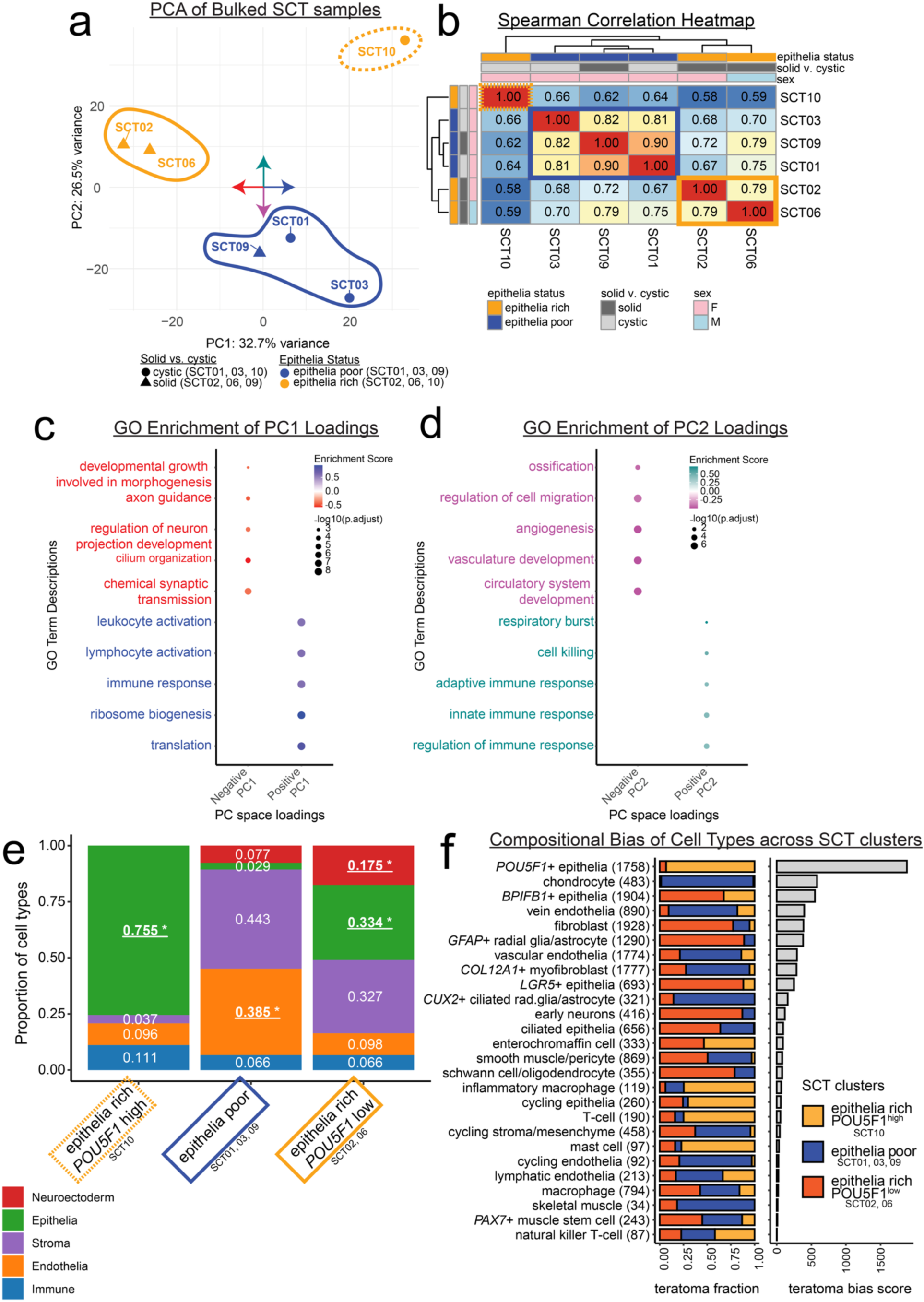
SCT transcriptional profiles cluster based on compositional differences. (a) Principal component analysis (PCA) plot of the snRNA-seq samples of SCTs with 3 clusters denoted by circles, *n*=6 SCTs. The cystic samples are mapped with a circle and solid samples are mapped with a triangle; colored bubbles: solid-orange is epithelia-rich *POU5F1*^low^ (SCT02/06), dashed-orange is the epithelia-rich *POU5F1*^high^ (SCT10), blue is epithelia-poor (SCT01/03/09). The compass in the center of the PCA plot denotes the gene loadings used for gene ontology (GO) analyses graphed in c. (b) Spearman correlation reveals that of the 3 clusters of SCT samples, SCT10 correlates least from the rest of the samples (dashed-orange box, epithelia-rich *POU5F1*^high^). In the center, the solid-blue box denotes the epithelia-poor samples which correlate strongest to one another. In the bottom left is the solid-orange box of epithelia-rich *POU5F1*^low^. Clustering was done by Euclidean distance. (c,d) PCA gene loadings for PC1 (c) and PC2 (d) were used in gene set enrichment analysis (GSEA) through gene ontologies within biological processes. Plotted are representative biological process terms for which there is enrichment in the top ranked genes for each loading, color coded to the PC space: red = negative PC1 score, blue = positive PC1 score, magenta = negative PC2 score, cyan = positive PC2 score. Benjamini-Hochberg corrected log-transformed p-values shown. (e) Bar plot colored by broad category cell identities, highlighting the significantly enriched epithelia population in epithelia-rich *POU5F1*^high^, the significantly enriched endothelial population in epithelia-poor, and the significantly enriched epithelia and neuroectoderm populations in Epithelia-rich *POU5F1*^low^ SCTs. (*) means a false discovery rate < 0.05 and absolute value of log_2_fold change > 1, the statistical test was a Monte-Carlo/permutation test. *n*=18,034 cells with 10,000 permutations (f) On the left: the proportional makeup of each sample across cell type, with total cell number in parentheses. On the right: the bias score for each cell type shows bias towards specific teratomas. A low bias score means the cell type is well mixed across the 3 clusters, whereas a high bias score means that the identity is made up by mostly one SCT group.

To understand the transcriptional programs driving this separation, we examined gene loadings for each PC direction (positive vs. negative) and conducted gene set enrichment analyses (GSEA) using Gene Ontology (GO) biological process terms ^41^. For PC1, which explained ∼33% of the variance, genes with negative loadings were enriched for neuronal and ciliary gene sets, while genes with positive loadings were enriched for translation machinery (including ribosomal functions) and immune activation gene sets (**Fig. 2c**, **Supplementary Table 3a,b**). PC2, which explained ∼27% of the variance, separated tumors based on distinct mesodermal programs: genes with negative loadings were enriched for bone formation and vascular development terms, while genes with positive loadings were enriched for immune response terms/pathways (**Fig. 2d** and **Supplementary Table 3c,d**). These results demonstrated that inflammatory, vascular, and neuroectodermal programs drive the major axes of transcriptional variation across SCTs.

Given that teratomas are thought to arise from a pluripotent progenitor and evolve through stochastic differentiation, we hypothesized that differences in cellular composition underlie the observed transcriptomic clusters ^19^. We therefore stratified samples according to their PC groupings and investigated their cell type distributions. SCT10 exhibited a distinct phenotype, composed almost entirely of *POU5F1*+ epithelial cells and entirely lacking neuroectoderm. SCT02, SCT06, and SCT10 shared a distinct epithelial population absent from SCT01, SCT03, and SCT09 (**Supplementary Fig. 2a**). Based on these findings, we defined three compositionally distinct tumor groups: (i) epithelia-rich *POU5F1*^high^ (SCT10) (ii) epithelia-poor (SCT01, SCT03, SCT09), and (ii) epithelia-rich *POU5F1*^low^ (SCT02, SCT06).

To investigate whether specific cell types were enriched in each stratified group, we statistically compared cell type proportions across groups. The epithelia-rich *POU5F1*^high^ SCT was significantly enriched for epithelial cells, the epithelia-poor SCTs were significantly enriched in endothelial cells, and the epithelia-rich *POU5F1*^low^ SCTs were significantly enriched in both epithelial and neuroectodermal lineages (False Discovery Rate < 0.05 and log-fold change > 1; **Fig. 2e, Supplementary Fig. 2b.i-ii**). To quantify the distribution of cell identities across tumors, we calculated a teratoma bias score, with lower scores indicating a more uniform distribution of a given cell type across samples (**Supplementary Fig. 2c**). Epithelial subsets (*POU5F1*+ and *BPIFB1*+ types), chondrocytes, venous and vascular endothelial cells, general fibroblasts, and radial glia/astrocyte populations (*GFAP*+ and *CUX2*+ types) showed the highest degree of bias, reflecting substantial inter-tumor variability.

Collectively, these results illustrate that PC space reflects distinct underlying cellular architectures. The epithelia-rich *POU5F1*^high^ sample (SCT10) was defined by transcriptional signatures related to immune activation and cell killing, and was enriched for inflammatory macrophages and T-cells. The epithelia-poor samples (SCT01, SCT03, SCT09) were driven by vascular development and ossification, and enriched for endothelial and chondrocyte populations, consistent with endochondral ossification. The epithelia-rich *POU5F1*^low^ samples (SCT02, SCT06) were characterized by neuronal development pathways and enriched for neuroectodermal cell types, specifically radial glia and astrocytes. Our findings provide a refined molecular framework for understanding inter-tumor heterogeneity in SCTs.

### Prenatally-resected sacrococcygeal teratomas are enriched for cycling stromal and neuroectodermal cells

SCTs lead to fetal demise when tumors are large, solid, immature, and rapidly growing, causing life-threatening complications such as cardiac failure ^7^. These high-risk cases often require *in utero* resection to prevent fetal loss or minimize morbidity (prenatally-resected SCTs, prSCTs). To characterize the cellular composition of these tumors, we performed SpTr on two FFPE-banked prSCTs (SCT04 and SCT05) (**Fig. 3a**, **Supplementary Fig. 3a**). Due to initial slide misalignment, two spatial datasets were generated for SCT04 (one from the tumor edge and one from the center), which showed high concordance (Spearman ρ = 0.99; **Supplementary Fig. 3b**) and were analyzed together. Visium SpTr 55 μm spots clustered by spatial locations (**Fig. 3b**, **Supplementary Fig. 3c,d**). Both prSCTs mapped closely in PC space to the epithelia-rich POU5F1^low^ cluster from the snRNA-seq atlas (**Fig. 3c**), indicating compositional similarity.

**Fig. 3.**
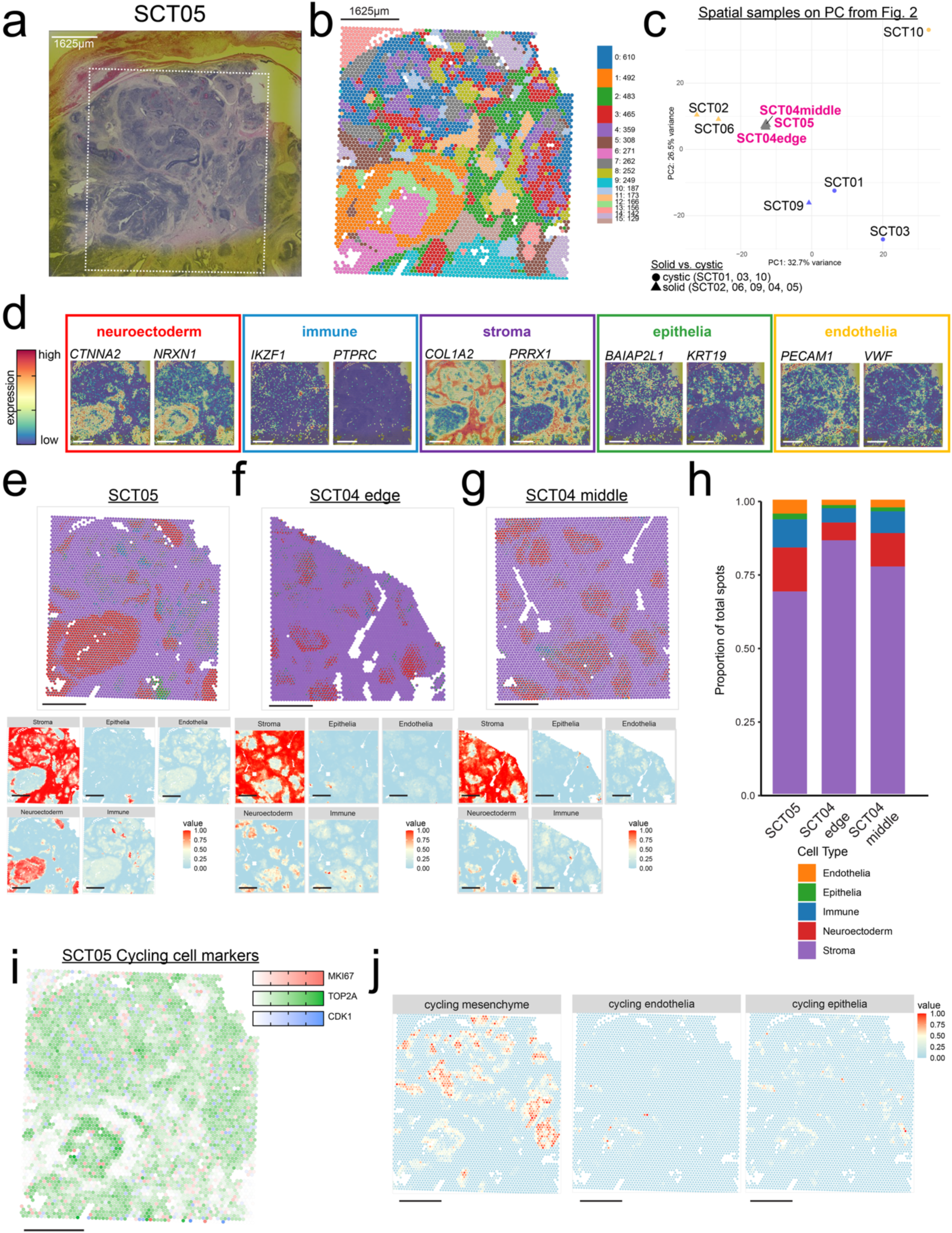
Spatial transcriptomics reveal composition of prenatal SCTs resembles post-natal SCTs (prSCT). (a) Hematoxylin and eosin staining of prSCT, SCT05, with box showing what area was used for spatial transcriptomics. (b) Unbiased clusters overlaid in spatial plot. Cluster numbered from largest to smallest, with the number of cells present in each cluster denoted. (c) Pseudobulked principal component analysis (PCA) plot from Fig. 2a with the spatial pseudobulked samples plotted (in magenta) using the same embeddings and genes. (d) Overlay of broad category markers taken from snRNA-seq atlas. Boxes correspond to the colors from Fig. 1c, neuroectoderm in red, immune in blue, stroma in purple, epithelia in green, and endothelia in orange. (e,f,g) Deconvolved spatial dots are shown in top graph as scatter pie chart, where each dot is a pie chart with the broad category proportion colored. SCT05 in e, SCT04 edge in f, SCT04 middle in g. On bottom is the spatial distribution of broad category intensities by deconvolution analyses. (h) Bar plot showing the proportional makeup of prSCTs colored by the corresponding cell type from snRNA-seq atlas. (i) Cycling cell genes expressed across the SCT sample, Colors correspond to each gene: red = *MKI67*, green = *TOP2A*, blue = *CDK1,* with white being no expression. (j) Spatial distribution of cycling cell subtype spatial intensities by deconvolution analyses. All scale bars = 1625µm.

To annotate broad cell types (**Fig. 2e**), we mapped expression of top marker genes identified from snRNA-seq on the prSCTs (**Supplementary Fig. 3e**). We observed high expression of stromal (*COL1A2*, *PRRX1*) and neuroectodermal (*CTNNA2*, *NRXN1*) markers across extensive regions of the prSCTs. In contrast, we observed more dispersed expression patterns in immune (*IKZF1*, *PTPRC*), epithelial (*BAIAP2L1*, *KRT19*), and endothelial (*PECAM1*, *VWF*) markers (**Fig. 3d**, **Supplementary Fig. 3f**). Visium SpTr spots encompass multiple cells, therefore we leveraged the CARD spatial deconvolution package ^42^ with our matched snRNA-seq atlas as a reference to estimate cell type composition at each spot (**Fig. 3e-g**). The prSCTs showed similar compositions overall, with the majority of spots comprising stromal (>60%), neuroectodermal (∼12%), and immune (<10%) populations (**Fig. 3h**), consistent with our marker gene expression patterns. These results suggest that prSCTs contain similar broad cell types as seen in the fresh SCT snRNA-seq atlas.

Given the rapid growth of these tumors, we sought to characterize cycling cell populations in the prSCTs using spot deconvolution. Among the cycling cells (**Fig. 3i**, **Supplementary Fig. 3g,i**), the majority contained *TOP2A*+ mesenchymal cells, with fewer spatial spots corresponding to cycling endothelial and epithelial populations (**Fig. 3j**, **Supplementary Fig. 3h,j**). In summary, SpTr of prSCTs revealed a predominant stromal and neuroectodermal composition, along with appreciable populations of cycling cells, providing insight into the cellular and transcriptional landscape of these rapidly growing prenatal SCTs.

### Profiled sacrococcygeal teratomas lack a distinct pluripotent or germ cell population

Given their histologic diversity, SCTs are thought to arise from a pluripotent progenitor: likely a migrating primordial germ cell (PGC) or a pluripotent PGC derivative ^9^. However, prior studies have not identified cells co-expressing canonical PGC or pluripotency transcription factors. We sought to identify a discrete cluster of single cells co-expressing both core pluripotency transcription factors and PGC markers (**Fig. 4a**, **Supplementary Fig. 4a**) ^43,44^. We did not observe a population co-expressing the core pluripotency transcription factors (*POU5F1*, *SOX2*, *KLF4* and *LIN28*) in any SCT. Likewise absent or sparsely expressed were early-stage PGC markers (pre-migratory; e.g., *PRDM1*, *TFAP2C*, *DND1*, *SOX17*) and late-stage germ cell markers (post-genital ridge colonization; *DDX4*, *DAZL*). While rare cells expressed one or two of these markers, none exhibited a full pluripotent or germ cell transcriptional signature (**Supplementary Fig. 4a**). Thus, although individual genes of interest were expressed sporadically, they did not constitute a bona fide pluripotent or germ cell-like population capable of initiating or propagating tumor differentiation and growth.

**Fig. 4.**
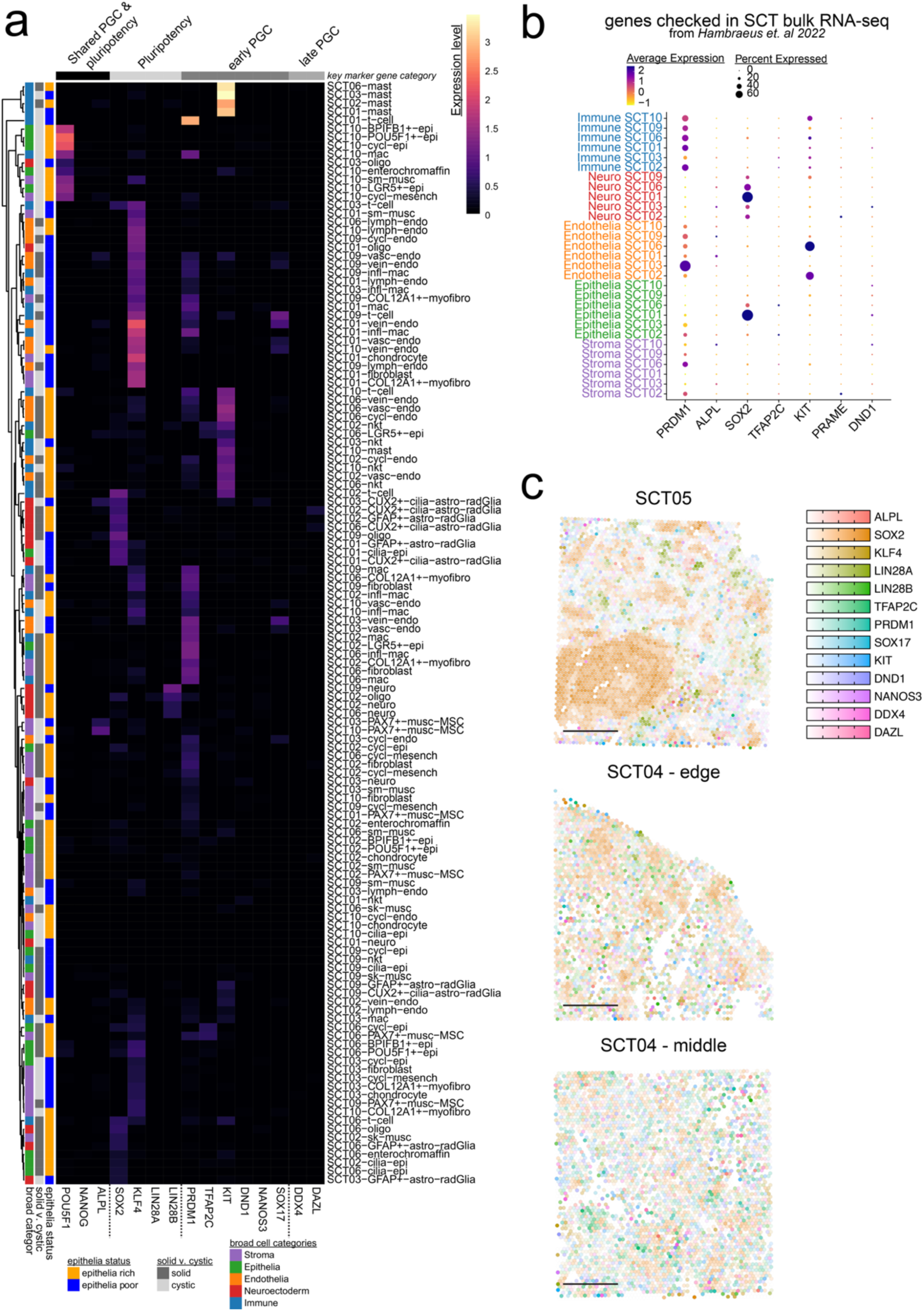
Primordial germ cell (PGC) and pluripotent genes do not co-occur in nuclei or spatial datasets. (a) Heatmap of the key genes in pluripotent cells and primordial germ cells (PGCs) grouped by the SCT sample and the cell types across the atlas. Column colors are the categories the genes belong to: black = genes shared in both pluripotent cells and PGCs, light grey = pluripotent only genes, dark grey = early PGC genes, medium grey = late PGC genes. Rows are clustered by Euclidean distances among the genes expressed. Row colors are the epithelial status (yellow = rich, blue = poor), solid vs. cystic categories (solid = dark grey, cystic = light grey), and the broad cell types (stroma = purple, epithelia = green, endothelia = orange, neuroectoderm = red, immune = blue). (b) DotPlot showing the average normalized expression and the percent of the cells expressing genes that were described as expressed in bulk RNA-seq of SCTs. (c) Pluripotent and PGC genes mapped onto the spatial dataset with no dots expressing all markers.

Previous bulk RNA-seq studies of SCTs had reported expression of genes typically associated with pluripotency maintenance (*ALPL* and *SOX2*) and early germ-cell specification (*PRDM1*, *TFAP2C* and *KIT*) ^14^. However, our snRNA-seq and SpTr analysis revealed that these genes were not restricted to a PGC- or pluripotent-like cluster, but rather distributed across differentiated cell types (**Fig. 4b,c**). Specifically, *PRDM1* expression was enriched in immune and endothelial cells; *TFAP2C* in immune and epithelial cells; and *KIT* predominantly in endothelial and mast cells ^45^. *ALPL* expression was detected in a small subset of endothelial cells, while *SOX2* was expressed in epithelial and neuroectodermal cells. Late fetal germ-cell markers, such as *PRAME* and *DND1*, were not robustly expressed in SCTs ^44^. In corroboration, SpTr profiling of prSCTs agreed with the snRNA-seq findings, revealing similar scattered expression of early germline genes without evidence of a defined germline-like niche (**Fig. 4c**). Together, these data demonstrate that SCTs do not harbor a molecularly defined population resembling pluripotent stem cells or PGCs. Instead, expression of select germline and pluripotency-associated genes occurs within differentiated tumor compartments, likely reflecting residual developmental gene expression rather than persistence of a progenitor state.

### Heterogeneous X-chromosome inactivation underlies cellular diversity in female sacrococcygeal teratomas

SCTs occur predominantly in individuals with two X-chromosomes (Xs), but the basis of this asymmetry remains unclear. X-chromosomal dosage compensation between sexes is achieved through X-chromosome inactivation (XCI) in XX cells, in which one X is silenced through a cascade triggered by the long non-coding RNA (lncRNA) *XIST* while the other X remains active (Xa). *XIST* coats the inactive X (Xi) in most somatic cells and mature PGCs ^10,46^. Given the female predominance of SCTs and the frequent loss of the Xi in cancers ^47^, we examined *XIST* expression across our SCT samples. We observed three distinct XIST expression patterns: 1) complete absence of *XIST*, consistent with an XY karyotype (SCT06); 2) ubiquitous *XIST* expression across all cell types (SCT01, 03, 09, 10); and 3) cell-type-specific *XIST* expression, predominantly in endothelial and immune compartments, but absent in others (SCT02) (**Fig. 5a,b**).

**Fig. 5.**
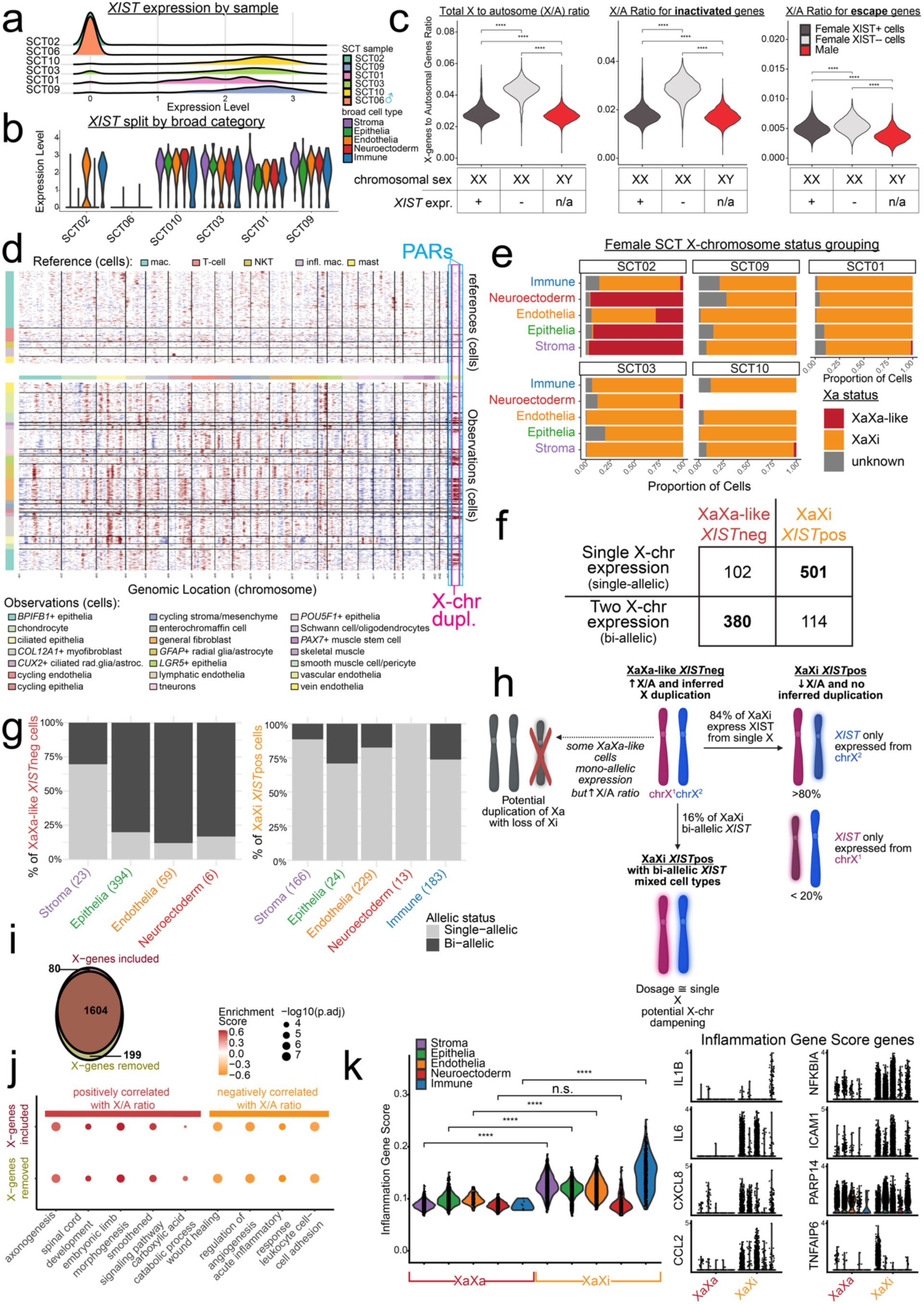
Profiling of SCT X-chromosome inactivation. (a) Ridgeline plot of *XIST* expression of SCT samples showing SCT02 and SCT06 have cells not expressing *XIST* (*XIST*neg). (b) Violin plot of *XIST* expression split by broad cell types and grouped by SCT samples showing that SCT02 endothelial and immune cells express *XIST* at comparable levels to the other XX SCTs. (c) X-to-Autosome (X/A) ratio violin plots stratified by XX cells that express *XIST* (*XIST*pos; dark grey), XX cells that do not express *XIST* (light grey), and XY cells (red). The three plots are then separated by overall X/A ratio, X/A ratio for X-linked genes that are inactivated during X-chromosome inactivation (XCI), and X/A ratio for X-linked genes that escape XCI. Wilcoxon ranked sum test: *n*=18,034 cells, **** = p-value < 0.0001. (d) InferCNV heatmap displaying inferred loss of a region as blue areas and inferred gain of a region as red areas. The pink box is the X-chromosome showing populations of cells that have a duplication event. The light blue boxes are the pseudoautosomal regions (PARs) of the X-chromosome. (e) Bar plots of the proportion of cell types that likely contain two active X-chromosomes (XaXa-like, burgundy) versus those that have one active X-chromosome (XaXi, orange). (f) Allele-specific RNA-sequencing of X-linked genes showing single-allelic vs bi-alletic expression in XaXa-like *XIST*neg (burgundy) and XaXi *XIST*pos (orange) cells. Bolding highlights that in the majority of XaXa-like cells, X-linked inactive genes were expressed bi-allelically, whereas in XaXi cells these genes were single-allelically expressed (*n*=1,097 cells, p-value < 0.0001, Chi-square with Yates’ correction). (g) Stacked bar plot showing single-allelic (light grey) and bi-allelic (dark grey) expression of inactive X-linked genes from allele specific RNA-seq. Top graph shows XaXa-like *XIST*neg cells and bottom shows XaXi *XIST*pos. Along the x-axis are cell types and number of cells in parentheses. (h) Diagram showing the hypothesized distribution of XCI differences among XaXi and XaXa-like cells. Figures made in Biorender. (i) Gene set enrichment analysis of genes correlated to the X/A ratio reveals biological process terms overlap whether the X-linked genes were included or removed. (j) Representative biological process terms plotted shows positively correlated terms in burgundy and negatively correlated terms in orange. (k) Violin plot inflammatory gene set score. A subset of genes from the gene set score is shown on the right. Wilcoxon ranked sum test: ****= p-value < 0.0001; n.s. = not significant, p-value > 0.05.

To investigate whether reduced XIST expression was associated with disrupted XCI, we calculated the ratio of X-linked to autosomal (X/A ratio) gene expression —a proxy for XCI status. In XaXi cells, the X/A ratio is approximately equal to XY cells, whereas in XaXa cells, the ratio is nearly double XY cells ^48,49^. Consistent with this pattern, XX cells expressing *XIST* (*XIST*pos) in SCTs had an X/A ratio similar to XY cells (0.029 vs. 0.027, p < 0.0001), whereas *XIST* negative (*XIST*neg) XX cells showed a nearly twofold increase (0.043 vs. 0.027, p < 0.0001) (**Fig. 5c**, **Supplementary Fig. 5a**). In SCT02, elevated X/A ratios were observed in all broad cell types except immune and endothelial cells, which retained high *XIST* expression (**Supplementary Fig. 5b**).

To further ascertain XCI fidelity, we stratified X-linked genes based on their known silencing status. There are two major categories of genes on the Xi, 392 inactivated genes, which are silenced by XCI therefore expressed only from Xa, and 82 escapee genes, which are not silenced therefore expressed from both Xa and Xi ^50^. To further investigate XCI variability in tumor cells, we computed the X/A ratio of inactivated and escapee genes (**Fig. 5c**). Among the 359 inactivated genes in our data, X/A ratios were similar between XY and *XIST*pos XX cells but nearly doubled in *XIST*neg XX cells (*XIST*pos XX = 0.018, *XIST*neg XX = 0.028, XY = 0.017, p < 0.0001). By contrast, expression of the 70 escapee genes in our data remained constant regardless of *XIST* status or sex (*XIST*pos XX = 0.005, *XIST*neg XX = 0.005, XY = 0.003, p < 0.0001) (**Fig. 5c**). This ratio remained similar across SCT samples as well (**Supplementary Fig. 5a**). These findings suggest that stratification of cells by *XIST* expression follows expected X/A ratios for X-linked genes in a normal XX karyotype.

To further characterize the Xs in the *XIST*neg cells, we compared the distribution of gene expression intensities across large segments of the X in our SCTs. Using inferCNV to calculate inferred copy number alterations, we identified areas of “duplications” and “deletions” on the X ^51^. SCT02 displayed the highest frequency of inferred duplications, spanning most of the X except the pseudoautosomal regions (PARs) (**Fig. 5d**, **Supplementary Fig. 5c**). The X/A ratio was positively correlated with the proportion of inferred X duplications (Spearman: 0.84; p < 0.0001) (**Supplementary Fig. 5d**). This analysis revealed two distinct cell clusters based on X/A ratio (>0.036) and inferred X duplication (>0.5). Cells above both thresholds were classified as XaXa-like and consisted of SCT02 epithelial, stromal, and neuroectodermal cells (**Fig. 5e**), with little or no *XIST* expression (**Fig. 5b**). While it is possible that *XIST* was not detected in these cells due to drop-out in scRNA-seq, the observed cell-type bias, inferred duplications, and elevated X/A ratio suggests that a subset of SCT02 cells exhibit XCI failure or reactivation of the Xi.

We next examined whether *XIST* expression in these tumors followed normal monoallelic behavior (**Supplementary Fig. 5d**). In cells with low X/A ratio and CNV duplications (XaXi), we assessed allele-specific *XIST* expression by genotyping single nucleotide polymorphisms (SNPs) on single-nuclei RNA molecules. If *XIST* reads are bi-allelically expressed—displaying variants with two alleles in individual cells—this result suggests cells are undergoing a process called X-chromosome dampening (XCD). In human preimplantation embryos and PGCs, XCD allows cells to achieve dosage compensation equivalent to a single X while expressing genes from both Xs ^52^. Conversely, if *XIST* is single-allelically expressed—showing variants from a single allele per cell — this would indicate canonical XCI rather than XCD. We found that *XIST* was predominantly expressed from a single X, consistent with largely normal XCI (**Supplementary Fig. 5e**). Within a tumor sample, most single chromosome reads were from the same X, implying that XaXi cells generally shared a common Xi. Taken together this implied that XCI is stable during tumor growth and potentially that SCTs may come from a clonal origin with a shared inactivated X (**Supplementary Fig. 5f**).

To distinguish between XCI failure and duplication of the Xa, we examined allele-specific expression of XCI inactivated genes. If XaXa-like cells exhibit bi-allelic expression of these genes it would indicate reactivation of the Xi or failure to undergo XCI, whereas single-allelic expression indicates potential loss of the Xi and duplication of the Xa ^53^ (**Fig. 5h**). SNP information can distinguish between these possibilities by asking whether X-linked gene expression arose from one or two Xs in XaXa-like *XIST*neg cells. These cells showed significantly higher rates of bi-allelic expression (79% bi-allelic vs 21% single-allelic), compared to XaXi *XIST*pos cells (19% bi-allelic vs. 81% single-allelic) (p <0.0001, **Fig. 5f**). Stratification by cell type revealed that XaXa-like *XIST*neg epithelia, endothelia, and neuroectoderm displayed bi-allelic expression, while 70% of stromal cells had single-allelic expression. Among the XaXi *XIST*pos cells, all lineages except stroma had single-allelic expression (**Fig. 5g**). Taken together, we find evidence that most XaXa-like cells have a second active X rather than a duplicated Xa with loss of Xi.

Finally, we explored the broader transcriptional consequences of increased X-linked gene dosage. Across all genes, we observed X/A ratio negatively or positively correlated with expression level of X-linked genes (**Supplementary Fig. 5g, Supplementary Table 4a**). GSEA of ranked Spearman correlations, performed with and without X-linked genes, showed 85% overlap in significantly enriched GO terms (**Fig. 5i, Supplementary Table 4b,c**). Genes positively correlated with X/A ratio were enriched for pathways including carboxylic acid catabolism, neuronal processes, Smoothened signaling, and embryonic limb development. In contrast, negatively correlated genes were enriched for immune and angiogenesis-related pathways (**Fig. 5j**). Comparing XaXa-like and XaXi cells revealed that inflammatory processes were significantly more active in XaXi cells across stromal, epithelial, endothelial, and immune cells, but not neuroectoderm (p < 0.0001; **Fig. 5k**). This data suggests that XCI failure contributes to cell-type-specific transcriptional shifts in SCTs, with broad implications for tumor phenotype and sex bias.

## Discussion

This study provides a comprehensive, single-cell atlas of SCTs, revealing detailed molecular and cellular stratification, highlighting inter-tumor compositional differences, and variation in XCI amongst SCTs. We identified five major lineages—neuroectodermal, epithelial, stromal, endothelial, and immune—each comprising multiple cell subtypes. SCTs clustered according to their cellular composition, suggesting a previously unrecognized distinction between epithelia-rich and epithelia-poor SCTs. Notably, we did not find evidence of a PGC or pluripotent progenitor in any sample. Additionally, we discovered XCI failure in a stromal, epithelial, and neuroectodermal cells in one tumor, associated with altered developmental and immune transcriptional signatures. Together, our findings uncover key cellular features of SCTs, offering new insights into tumor heterogeneity and potential avenues for improved diagnostics and clinical stratification.

A key discovery was the stratification of SCTs into epithelia-rich and epithelia-poor subtypes based on lineage composition. Epithelia-rich tumors were enriched for neuronal and epithelial markers, while epithelia-poor tumors exhibited strong immune and vascular signatures, suggesting divergent differentiation trajectories and tumor microenvironment influences. Clinically, SCTs are classified by histologic composition ^2,4,6^, but our study refines this framework by identifying transcriptomic signatures that more precisely define these subtypes. Notably, spatial profiling of prSCTs, the most aggressive tumor in our cohort, revealed that these tumors most closely resemble the epithelia-rich class in PC space, suggesting that compositional state may relate to clinical behavior. The stratification lays the groundwork for the development of multiplexed antibody-based histopathology, offering a more precise and informative diagnostic approach.

Our characterization of prSCTs with SpTr showed extensive expression of cycling cell markers, consistent with their rapid *in utero* growth. We identified cycling populations of stroma, endothelia, and epithelia, reminiscent of transient amplifying cells– the source of tumor growth within growing teratoma syndrome, a rare complication of mature teratomas involving expansion of non-malignant tissue during or after chemotherapy ^54^. The presence of cycling populations in stromal, endothelial, and epithelial cells within prSCTs and our broader snRNA-seq atlas suggests that a transient amplifying population may drive SCT expansion. Moreover, the enrichment of neuroectoderm components in prSCTs echoes findings in ovarian teratomas, where primitive neuroectoderm content is prognostic^55^; this finding raises the possibility that similar features could serve as potential prognostic markers, warranting further investigation.

Given the female predominance of SCTs, we investigated XCI as a potential contributor to SCT biology. We identified a population of XaXa-like cells, characterized by high X/A ratios, inferred X-chromosome duplications, and loss of *XIST* expression, primarily within SCT02 epithelia, stroma, and neuroectoderm. The XaXa-like cells express low levels of inflammatory genes compared to the same lineages in the XaXi cells. XaXa-like cells also showed biallelic expression of normally inactivated X-genes, consistent with XCI failure or X reactivation^56^. However, some stromal cells retained monoallelic expression despite apparent X duplication, possibly reflecting Xa duplication with Xi loss^53^. Endothelial and immune cells in SCT02 largely maintained XaXi status, supporting prior findings that these populations are infiltrative and not intrinsic to the tumor^18^.

Differences in X-chromosome activity within SCTs could suggest that XCI may not be uniformly stable across tumor cells. However, our SNP-based analysis of *XIST* RNA reads within XaXi cells revealed cells within the same tumor expressed *XIST* from a shared X across multiple lineages, supporting a model in which these cells are clonally derived from a common precursor that had already undergone XCI. This finding is consistent with prior reports of skewed XCI patterns in SCTs ^57^, and provides additional evidence that X-linked activity gene dosage may be dysregulated in a lineage-specific manner. Together, these results introduce potential mechanisms for X-linked gene dysregulation in SCTs. Such dysregulation may help explain the strong sex bias in SCT incidence and warrants further investigation^58^.

Although previous studies reported the expression of pluripotency and germ cell-related proteins in SCTs, our single-nuclei or spatial data did not reveal a bona fide PGC or pluripotent population^13^ ^12^. Our atlas revealed that these genes are diffusely expressed across multiple cell types rather than coincident in both snRNA-seq and SpTr. For example, *POU5F1*+ epithelial cells expressed few pluripotency genes and resembled regenerative lung epithelia ^59,60^. While the role of *POU5F1* in SCTs remains unclear, its recurrent expression in other cancers invites the possibility that it may contribute to tumorigenesis through mechanisms unrelated to germ cell identity ^61^.

By combining single-cell and spatial methods, we dissected SCT heterogeneity, enabling intra- and inter-tumor comparisons. Unlike previous studies that relied on hESCs in immunodeficient mice, our atlas offers insights into development within undirected differentiation in a clinical disease. Importantly, we show that spatial context is essential for interpreting SCT composition, as single-region sampling may miss intra-tumoral heterogeneity. Future studies should prioritize multi-region profiling to capture this complexity more comprehensively.

We acknowledge the presence of limitations in our study. Fresh tumor samples were not perfused prior to dissociation, potentially confounding the distinction between intravascular and extravascular immune cells. Nonetheless, spatial data support the interpretation that these immune cells are tumor-infiltrating. Due to the rarity of this tumor type, our sample size is small, which affects the generalizability of this study to all SCTs. Additionally, SpTr lacks probes for lncRNAs, such as *XIST*, limiting direct comparison with snRNA-seq. The allelic SNP analyses also suffer from limited coverage of each X-chromosome gene in our snRNA-seq data due to the constraints of 10X Genomics chemistry.

Looking ahead, our clinical pathology banked samples offers an opportunity to assess recurrence risk by comparing primary and recurrent SCTs. Incorporating FFPE-compatible snRNA-seq techniques with expanded probe sets for *XIST* and other lncRNAs will enhance analysis of archival specimens^62^. Investigating XCI and X-linked gene dosage in larger, diverse SCT-cohorts may uncover new regulatory mechanisms and therapeutic targets. Future probe-based FFPE snRNA-seq studies should include probes for XCI lncRNAs to permit assessments of cell-type specific XCI dysregulation, as identified in this study. Deeper allelic SNP analyses and spatially resolved sampling will be essential for linking chromosomal architecture to functional outcomes.

The identification of XaXa-like cells with XCI failure has broad implications for tumor biology. Increased X-linked gene dosage could drive transcriptional imbalance, accelerate tumor progression, or confer resistance to therapy, particularly in cell types where X-chromosome dosage-sensitive pathways are affected (e.g., immune suppression, developmental signaling). Mechanistically, XCI defects in tumors could arise from several processes, including epigenetic instability, developmental mis-programming, or selective pressures that favor cells with elevated X-linked expression. These findings suggest that XCI heterogeneity may not be a byproduct of tumor evolution, but a contributor to disease trajectory. Therefore, uncovering the drivers and consequences of XCI dysregulation in SCTs, and other germ cell tumors, may ultimately yield new biomarkers and interventions.

## Materials and Methods

### Patient selection and inclusion for SCT samples

To investigate intratumor heterogeneity in sacrococcygeal teratomas (SCTs), we collected eight fresh SCT samples from patients aged 1 day to 4 months at UCSF and Colorado Children’s Hospital, along with two banked fresh frozen paraffin-embedded (FFPE) samples from UCSF. We selected only sacrococcygeal teratomas for sequencing, excluding one cervical and one pericardial teratoma. Informed consent was obtained from each patient via their parent or legal guardian. The UCSF and Colorado Children’s Hospital IRBs approved the collections (UCSF IRB#: 10-00093; Colorado Children’s Hospital IRB#: 22-1906).

For the banked samples at UCSF used in spatial transcriptomics, we searched the internal pathology database for sacrococcygeal teratoma cases from 2011-2016 and selected those with adequate tissue. Informed consent was waived based on the IRB’s criteria that the research posed minimal risk, did not adversely affect subjects’ rights and welfare, was impracticable without the waiver, and additional pertinent information would be provided when appropriate (UCSF IRB#: 16-19356).

### Teratoma isolation from surgical resection, freezing, and thawing

Teratoma pieces (< 0.5 cm³) were taken from the solid component of the tumor for all samples. For SCT01, only the cystic wall was taken due to its predominantly cystic nature. Except SCT01, all samples were frozen in CryoStor 10 (CAT#07952, StemCell Technologies) overnight in freezing containers that cool at approximately -1°C/minute. SCT01 was frozen in 10% DMSO in FBS and stored at -80°C due to technical limitations. SCT01 yielded 253 cells, likely due to its storage conditions. All samples were thawed in PBS without Ca/Mg until fully thawed. Samples were processed in pairs (SCT01 and SCT02; SCT03 and SCT06; SCT09 and SCT10) to have one solid and one cystic sample processed together in each batch.

### Teratoma dissociation of fresh sample for single cell RNA-sequencing

To isolate cells for single-cell RNA-seq (scRNA-seq), teratoma pieces (< 0.5 cm³) were thawed and dissected into smaller pieces using sterile scalpels in PBS without calcium and magnesium. The pieces were placed in 0.25% collagenase IV with 0.25% trypsin for 30 minutes with agitation every 10 minutes. After digestion, the suspension was filtered through a 100 µm filter. Live cells were sorted using Sytox Red on a BD FACSAria2 SORP (BD Biosciences, cat# 643245) and prepared for sequencing as written below.

### Teratoma dissociation of fresh sample for single nuclei RNA-sequencing

For single-nuclei RNA-seq (snRNA-seq), we used the 10X Genomics Chromium Nuclei Isolation Kit (CG000505 Rev. A). Teratoma pieces (< 0.5 cm³) were dissected into pieces (< 5 mg each) on a dry ice-cooled glass petri dish and placed in Sample Dissociation Tubes. No more than ∼50 mg of tissue was used, with 200 µL lysis buffer added before homogenization with a plastic pestle. After adding 300 µL lysis buffer, samples were incubated for 10 minutes, centrifuged, and the flow-through isolated. The nuclei pellet was resuspended in 500 µL Debris Removal buffer, centrifuged, and supernatant removed. The pellet was washed once in Wash & Resuspension Buffer and resuspended in < 200 µL, ready for counting.

### Preparing sequencing of single cell and nuclei from teratomas

Filtered cell and nuclei suspensions (40 µm and 100 µm filters, respectively) were counted using the Cellaca MX High-throughput Automated Cell Counter (CAT# MX-112-0127, Nexcelom) and adjusted to ∼1000 cells/µL in Wash and Resuspension Buffer or 0.04% BSA-PBS. Library preparation was performed using the 10X Genomics Chromium Next GEM Single Cell 3’ kit (CG000315 Rev.F) with dual-indexing following manufacturer instructions. Nuclei/cells were prepared for Gel Beads-in-emulsion (GEMs) with a final concentration capturing ∼10,000 cells per sample. 10X primers amplified polyA transcripts per cell, with enzymatic fragmentation and size selection optimizing cDNA amplicon size. Illumina-compatible libraries were quantified with a BioAnalyzer before loading onto a Novaseq6000 instrument targeting 20,000 reads per cell. Multiplexing several samples maximized data output.

### Spatial transcriptomic slide preparation with Visium

Sections from FFPE blocks stored at room temperature were cut onto charged glass slides at 14 µm thickness. RNA quality was tested by extracting RNA from two adjacent sections using the RNEasy FFPE kit (73504, Qiagen) and measuring DV200 scores with a BioAnalyzer and RNA Pico 6000 kit (5067-1513, Agilent). Only samples with DV200 scores >30% were used. Slides were prepared for spatial transcriptomics using the manufacturer’s Demonstrated Protocol (CG000520 Rev. B, 10X Genomics), including deparaffinization, hematoxylin and eosin staining, and imaging with a Keyence BZ-X series microscope at 20X magnification. Slides were then processed according to CG000520 and de-crosslinked.

### sc/snRNA-seq quality control and filtering

Demultiplexing of sample library reads was completed by the Chan Zuckerberg Biohub, and FASTQ files were then aligned using CellRanger-7.0.1 from 10X Genomics with the default human reference genome (refdata-gex-GRCh38-2020-A).

For all 10X libraries, initial quality control was performed separately for each sample. Counts matrices were corrected for ambient RNA removal using the SoupX package (v1.6.2)^63^. Briefly, raw and filtered CellRanger output count matrices were read into R (v4.3.0) using the Seurat package (v5.3.0) ^64^ with the Read10X function. A soup object was created from the raw and filtered count matrices, while the filtered matrix was used to create a Seurat object using the CreateSeuratObject function. The Seurat analysis pipeline (NormalizeData, FindVariableFeatures, ScaleData, RunPCA, FindNeighbors, FindClusters, RunUMAP) was executed using default parameters and 20 PCA dimensions to provide clustering and dimensionality reduction information for the soup object using setClusters and SetDR. Ambient RNA counts were identified and removed using the autoEstCont and adjustCounts functions.

New Seurat objects were created using the adjusted count matrices and had doublets removed. The scCustomize package (v3.0.1) ^65^ was used to visualize all analyses. The Add_Cell_QC_Metrics function was used to calculate mitochondrial and ribosomal percentages, cell complexity (log10GenesPerUMI), and top gene percentages (oxidative phosphorylation, apoptosis, DNA repair, and blood genes). Filtering was then performed to remove low-quality cells by applying a cutoff of at least 1500 reads per cell, at least 1000 genes per cell, and no more than 20% mitochondrial genes for scRNA-seq and 5% for snRNA-seq.

Doublets were removed using DoubletFinder (v2.0.4) ^66^ and scDblFinder (v1.16.0) ^67^. DoubletFinder identified potential doublets based on the rate of doublets expected in the dataset, involving parameter sweeps to identify the optimal pK and homotypic doublet proportion estimation, followed by the application of the DoubletFinder algorithm to classify cells as doublets or singlets. scDblFinder was then applied to further refine doublet identification by converting the Seurat objects to SingleCellExperiment objects ^68^ and running the scDblFinder algorithm to classify cells and nuclei. The classifications from both DoubletFinder and scDblFinder were used to retain only singlets for downstream analysis.

Low-quality singlet cells were removed based on RNA counts between 1500-25000 and gene counts above 1000. The mitochondrial gene cutoff used was 20% for single-cell RNA-seq and 5% for single-nuclei RNA-seq. Genes were filtered by keeping only those expressed in at least 10 cells. All mitochondrial genes were removed from single-nuclei samples as they were expected to be ambient reads. We had a total of 18,034 cells at the end.

### Dimensionality Reduction, Clustering, and Annotation of sn/scRNA-seq

With the filtered Seurat objects for each sample, we analyzed each sample separately. The objects were log-normalized using NormalizeData (normalization.method = “LogNormalize”, scale.factor = 10000), and the top 2,000 variable genes were selected, after which the data were scaled using the default ScaleData. We computed 60 principal components (PCs) and determined the optimal number for each sample using an ElbowPlot; these optimal PCs were then used to identify clusters at a coarse resolution of 0.1. Differentially expressed genes for each cluster were determined using the Wilcoxon Rank Sum test via the FindAllMarkers function (logfc.threshold = 0.25, only.pos = TRUE). Cell types in the split objects were annotated using four distinct approaches: first, automated cell annotation via a hierarchically organized marker gene map ^17^ by inputting the top 50 genes from each cluster into ACT and retaining the top annotation (clusters without a clear cell-type correlation were left unannotated). Second, we correlated each cluster with published ovarian (GEO: GSE229343) and human embryonic stem cell (ESC)-derived teratoma (GEO: GSE156170) datasets ^18^,^19^ generating transcriptional signatures for each reference cluster by selecting the top 100 differentially expressed genes (ranked by increasing adjusted p-value) and scoring the query dataset with Seurat’s AddModuleScore function. Third, we mapped our clusters onto a human fetal development atlas ^20^. using Seurat’s RunAzimuth, running the “fetusref” from Cao et al. (2020) and retaining the top average score for each cluster. Lastly, cell type classification was performed using a large language model-based framework ^21^. where the top 50 genes from each cluster were provided to ChatGPT’s gpt-4o model (May 24, 2024 version) along with the tissue name “human sacrococcygeal teratoma”. These results are summarized in Supplemental Table 1.

To integrate both (i) all nuclei samples and (ii) just the SCT01 cells and nuclei, we used the Seurat workflow by first splitting the RNA assay based on the individual dataset identifier with the split function; the count matrix was normalized and variable features were identified using default parameters. Feature counts were scaled with ScaleData, and PCA was performed using default parameters with optimal PC numbers chosen from an ElbowPlot. These PCs were then used for integration via the IntegrateLayers function with HarmonyIntegration, and a nearest neighbor graph along with UMAP dimensionality reduction was computed based on the Harmony reduction. Broad cell type clusters were identified using Louvain clustering, and their relationships across resolutions ranging from 0.1 to 1.2 (by intervals of 0.1) were visualized using a clustree with a Sugiyama layout ^69^. Finally, the integrated SCT samples were annotated at the highest resolution (0.7) using distinct cell names to yield 26 separate clusters (sub-categories), which were then compared to corresponding cell types in the split annotations and to the published ovarian and hESC-derived teratoma datasets ^18^,^19^. A combination of the top differentially expressed genes and known marker genes was used to assign cell type identities to each sub-category, as detailed in the Results.

### Pseudobulked principal component analysis

To evaluate global transcriptional patterns across SCT samples, we performed pseudobulked principal component analysis (PCA) using the RNA counts from the integrated snRNA-seq dataset. In order to control for variability in cell numbers across samples, we downsampled each sample to have an equal total number of cells (253 per sample since SCT01 had that many nuclei). Following this downsampling, we used Seurat to normalize the data (NormalizeData) and identified variable features (FindVariableFeatures). FindVariableFeatures was used in default mode, which uses the “vst” method to find the top 2000 variable genes. With a robust set of variable genes in hand, we aggregated expression values for each sample to create a pseudobulk expression matrix for each sample using Seurat’s AggregateExpression and only the highly variable genes were retained for downstream analyses.

We then constructed a DESeq2 dataset using the aggregated data and sample metadata, which included relevant biological covariates such as sex and tumor phenotype (solid or cystic). Genes with counts below 10 were filtered out to ensure sufficient data quality for downstream analysis, but no genes were lower than 10 counts. Given the limited number of samples, we implemented DESeq2 regularized log (rlog) transformation to stabilize variance across the dataset, ensuring that technical variation did not confound biological signals. PCA was then conducted using the rlog-transformed data, with the expression matrix being scaled and centered. The resulting principal components were used to quantify the percentage of explained variance per component.

We extracted the loadings for PC1 and PC2 from the PCA rotation matrix for downstream pathway enrichment analysis. These loadings represent the contribution of each variable (gene) to the corresponding principal component, allowing us to rank the genes that most strongly influence the observed variance. Specifically, we extracted the loadings for PC1 and PC2, sorting these values in descending order to highlight the top contributors to each direction. This process facilitated the identification of genes that drive the major transcriptional patterns captured by each PC.

### Differential proportional testing

Differential proportion testing on SCT samples was run using permutation testing with scProportionTest (v0.0.0.9) ^70^. Briefly, cell identities were randomly shuffled between the two samples, and the Log2 proportional difference between the cell counts for each population was computed. This process was repeated 5,000 times, and the p-values were calculated as the number of proportional differences that were as or more extreme than the observed value (plus one) divided by the total number of iterations (plus one) ^71^. These p-values were then adjusted for multiple comparisons using false discovery rate (FDR) correction. Bootstrap sampling was employed to generate 95% confidence intervals for plotting. In each iteration, cell identities for each sample were sampled with replacement, and the log2 fold difference between cell counts in each population was obtained. This procedure was repeated 5,000 times, and the confidence interval was defined as the range between the 2.5th and 97.5th percentiles of the simulated log2 proportional differences.

### Teratoma bias score

We followed the process as published by McDonald et. al. 2020 ^19^. Briefly, to quantify the heterogeneity between SCT samples and the Epithelia split, we employed the Normalized Relative Entropy metric from CONOS ^72^. For each cell type (k), we constructed a vector representing the number of cells in each sample and computed the empirical KL divergence between this vector and the overall cell distribution across all samples. A higher Normalized Relative Entropy indicates that the cell types are more evenly distributed (i.e., less biased), while a lower value suggests that a cell type is more concentrated in a specific sample (i.e., more biased). To further assess the bias of individual cell types, we calculated the KL divergence between the cell count distribution for each cell type and the overall distribution, then scaled this divergence by the total number of cells in that cell type.

### Sequencing for 10X Visium

Following slide preparation, the slides had libraries constructed using the manufacturer’s Demonstrated Protocol (CG000495 Rev. E, 10X Genomics). Briefly, the manufacturer probes were hybridized to the sample’s RNA then ligated. Then, the ligation products are released from the tissue and then captured on the Visium slides with an additional UMI barcode. Following these steps the libraries are amplified and prepared for sequencing. Sequencing was completed on a NextSeq 2000 instrument with a minimum of 25,000 read pairs per tissue covered spot on Capture Area as calculated by the percentage of tissue on the capture area. Multiplexing of several samples run together was completed to maximize the run of data.

### Spatial transcriptomics data alignment and pre-processing

Demultiplexing of sample library reads were completed by the Chan Zuckerberg Biohub, and fastq files were then aligned using spaceranger-2.1.0 software from 10X Genomics. The reference genome option (“--transcriptome”) used was the default human 10X reference genome, refdata-gex-GRCh38-2020-A. To create the Loupe Alignment Files (“--loupe-alignment”), the images taken from the CytAssist and the images taken on the Keyence BZ-X were imported into LoupeBrowser v7.0.1. The images were manually aligned, and the dots overlaying tissue were manually annotated as containing tissue. These data were all run on a High-Performance Computing Cluster at UCSF, C4, using an array-based script.

### Spatial transcriptomics processing and analysis

Comprehensive data analysis, including filtering, normalization, dimensionality reduction, and unsupervised hierarchical clustering, was performed using open-source R pipelines (Semla v1.1.6 and Seurat) ^73^ alongside custom in-house R code. Deconvolution spatial spot analysis was performed using CARD ^42^ and this publication’s SCT snRNA-seq dataset as reference. A total of 12,972 spatial spots were analyzed, covering on average 5400 genes distributed across 16 spatial clusters on average. Filtering criteria applied during preprocessing included a minimum of 50 UMI counts per gene, at least 5 spots per gene, and a minimum of 250 UMI counts per spot.

### Mapping spatial transcriptomics data to snRNA-seq PCA

To compare our spatial transcriptomic profiles with the dissociated nuclei dataset, we first aggregated spot-level counts for each spatial section to produce spatial pseudobulk profiles. These profiles were restricted to the same set of highly variable genes (HVGs, n = 2,000) that defined the snRNA-seq analysis, and the matrix was transformed with DESeq2 regularized-log function in parallel with the nuclei data.

The rlog-transformed spatial matrix was then projected onto the principal-component space previously defined by the snRNA-seq pseudobulk PCA using the predict() function on the existing principle component object (prcomp). This provided PC scores for each spatial sample on the identical axes. We visualized spatial and nuclei samples together (PC1 vs PC2), colouring by epithelia-rich/poor status and shaping by solid versus cystic phenotype, and calculated Euclidean distances between spatial and nuclei points to identify the closest transcriptional match for each spatial section.

### X-chromosome gene mapping and ratios

To find the X to Autosome (X/A) Ratio, we followed previously described methods ^49^. Briefly, we connected to the Ensembl database (GRCh38/hg38) to retrieve gene coordinate data for genes using biomaRt ^74^. Gene information, including HGNC symbols, chromosome names, start and end positions, and strand orientation, was obtained, and entries with empty gene names were excluded from subsequent analyses. For each cell, we calculated the total expression of X-chromosome genes and autosomal genes by summing their respective raw counts. The ratio of X-chromosome to autosomal gene expression was then computed by dividing the summed expression of X-chromosome genes by that of autosomal genes. X-chromosome genes were also stratified based on the X-chromosome inactivation (XCI) status into genes typically inactivated and those typically escaping XCI ^50^.

### Inference of copy number alterations

To infer copy number alterations (CNAs), we employed the inferCNV tool v1.19.1 ^51^. Briefly, inferCNV requires as input a filtered raw count matrix, a file specifying the order and chromosomal locations of genes, and a list of reference cells. In this study, we designated all immune cells (macrophages, inflammatory macrophages, T-cells, natural killer cells, and mast cells) as the reference population given their noted infiltrative patterns in teratomas ^18,19^. To generate the inferCNV object, we used the CreateInfercnvObject() function with default parameters (no maximum cells per group, minimum 100 counts per cell, and no mitochondrial genes). Subsequently, we ran the inferCNV function (infercnv::run()) with default settings (cutoff = 0.1, cluster_by_groups = TRUE, denoise = TRUE, and HMM = TRUE). Within this pipeline, genes with low expression levels across cells were first filtered out to reduce noise. The expression data were then normalized and log-transformed, and the mean expression of each gene (calculated from the reference cells) was subtracted from each cell to emphasize deviations from a normal state. Following normalization, a smoothing step that applies a moving average along each chromosome was performed to capture gradual shifts in expression. Finally, a Hidden Markov Model (HMM) was applied to predict CNV states (such as deletion, neutral, or amplification). The resulting output was visualized and integrated into the Seurat object using the infercnv::add_to_seurat function. The CNA data extracted through this process were subsequently used for mapping X-chromosome dynamics.

### Allele specific RNA-sequencing analyses

RNA variant calling was performed following the preprocessing workflow outlined by scLinaX ^75^. Briefly, aligned BAM files were filtered to retain only reads mapping to X-chromosome genes. These X-chromosome–specific BAM files were then processed using cellsnp-lite in mode 2a ^76^, with parameters set to a minimum count of 20 (--minCOUNT 20), a minimum minor allele frequency of 0.05 (--minMAF 0.05), and a minimum mapping quality of 20 (--minMAPQ 20). A custom bash script was employed to iterate over the SCT samples (e.g., “SCT01_nuc,” “SCT02,” “SCT03,” “SCT06,” “SCT09,” “SCT10”), using a predefined reference genome (GRCh38) and barcode files to generate summary SNP call files. Subsequently, gene-based annotation of the variants was conducted using Annovar ^77^. Although the original scLinaX protocol suggests incorporating allele frequency filtering (e.g., using the avsnp150 and ALL.sites.2015_08 databases), only gene-based annotations were applied here due to difficulties in sourcing the recommended references. The Annovar command was thus configured to utilize the refGene and ensGene databases for annotating gene locations. Finally, a modified version of the scLinaX quality control pipeline was implemented in R to merge the raw SNP call data with the Annovar annotations. This step involved generating a unique SNP identifier, filtering out variants with multiple gene assignments, and retaining only variants located within regions of interest (i.e., intergenic, intronic, UTR, exonic, ncRNA exonic, ncRNA intronic, and splicing regions). This workflow ensured robust allele-specific annotation of RNA variants for downstream analyses. XCI genes were taken from same methods in “X-chromosome gene mapping and ratios”.

### X-chromosome gene correlations

To evaluate the relationship between gene expression and X-chromosome dosage relative to autosomes, we computed Spearman correlation coefficients for each gene using the log-normalized expression data from the Seurat object and the corresponding X-to-Autosome ratio for each cell. Briefly, for each gene, we performed a Spearman correlation test (cor.test) to obtain the correlation coefficient and p-value while handling missing values using complete observations. To account for multiple comparisons, p-values were adjusted using the Benjamini-Hochberg FDR correction, and q-values were additionally computed using the qvalue package v2.34.0. The resulting data frame, containing gene identifiers, correlation estimates, original p-values, adjusted p-values, and q-values, was then sorted in descending order based on the absolute correlation coefficients to highlight the genes with the strongest associations in both positive and negative directions. The dataframe was also evaluated for gene locations using the Ensembl database (GRCh38/hg38) from X/A ratio to find what genes were autosomal vs. X-chromosome.

### Gene Set Scores with UCell

We used UCell to add gene set scores to our data as previously described ^78^. UCell was chosen for its robust performance across single-cell datasets, as it ranks genes within each cell rather than relying on absolute expression values—a feature that makes it resilient to variations in overall expression levels across datasets ^79^. Using UCell, we calculated gene set scores by providing a list of genes that define the signature of interest. For instance, to derive scores for hallmark gene sets, we retrieved the relevant gene lists from the Molecular Signatures Database (MSigDB) via the msigdbr package (v7.5.1) ^80^. For each cell, UCell first ranks all genes based on their expression levels. It then computes a Mann-Whitney U statistic for the provided gene set, which reflects how consistently the genes in the set are expressed at higher ranks relative to all other genes in that cell. The resulting UCell scores were integrated into our Seurat objects and subsequently visualized as violin plots using scCustomize, Seurat, and ggplot2.

### Pathway enrichment analysis

Pathway enrichment analysis was conducted using the clusterProfiler package (v4.10.1) with gene set enrichment analysis (GSEA) on Gene Ontology (GO) terms, facilitated by the gseGO function. We used gene list ranked by log-fold change, with gene symbols as key identifiers, adjusting p-values using the Benjamini-Hochberg (BH) method, and setting a p-value and q-value cutoff of 0.05. The human annotation database, org.Hs.eg.db (v3.18.0), was used for mapping gene symbols to GO terms. The analysis encompassed all GO categories, including biological process (BP), cellular component (CC), and molecular function (MF), with results from each category combined. Parameters were set to include gene sets with a minimum size of 10 and a maximum size of 500 genes, and 10,000 permutations were conducted to ensure robust statistical assessment. For visualization, we focused on BP terms that met significance criteria (adjusted p-value < 0.05, false discovery rate < 0.05) and had an enrichment score above 0.4 or below -0.4. These terms were further refined for semantic similarity using the simplify function ^81^. Treeplots were run with default parameters on the full GSEA results to visualize GO enriched themes. The curated terms were selected and reviewed by the authors for representation.

### Quantification and Statistical Testing

To determine statistics on the single-cell data for continuous variables, including gene expression, X/A ratio, and UCell scores, we used a non-parametric Wilcoxon rank-sum test in R. Given the low number of samples, technical replicates were not compared via pseudobulking approaches. To compare the number of XaXa-like cells and XaXi cells expressing bi-allelic versus mono-allelic X-linked genes, we performed a chi-square test with Yates correction to account for small sample sizes using GraphPad Prism. To assess the similarity of teratomas to each other, we utilized the pseudobulked matrices of each tumor to run Spearman correlation. Spearman correlations were computed on the rlog-transformed pseudobulked counts matrix and visualized using R. Among all analyses, significance was considered if corrected p-values were below 0.05.

### Declaration of generative AI and AI-assisted technologies in the writing process

During the preparation of this work the author(s) used UCSF VersaChat (ChatGPT version 4o) and OpenAI ChatGPT (version o3-mini-high) in order to (1) improve readability and brevity of the work the authors had written and (2) asking for feedback on the clarity of writing and the interpretation of the flow of logic (but not for experimental feedback). In accordance with the current Elsevier publishing ethics, after using these tools/services, the author(s) reviewed and edited the content as needed and take(s) full responsibility for the content of the published article.

## Data availability

The datasets generated during and/or analysed during the current study are available from the corresponding author on reasonable request. We are currently in the process of submitting raw files to dbGaP Study Submission. Meanwhile we are reviewing IRB approvals to also place all sequencing data and expression count matrices in Gene Expression Omnibus (GEO) and the cellxgene portal. Data will be publicly available prior to publication.

## Code availability

Code to produce all figures is available upon request. But will be available publicly prior to publication at: https://github.com/ejscience/2025_Rojas_SCTatlas.

## Acknowledgements

Most importantly, we thank the families who consented to donate their child’s SCT tissue for this study. We would like to thank the clinical team members from the MacKenzie and Derderian labs (E. Canepa, H. Isberg) for their help in establishing the IRBs and patient advocacy.

We would like to thank the teams and members of the UCSF CoLabs Initiative, a new model for research collaborations and core labs for the UCSF community. Specifically the Genomics CoLab (especially A. Poon, C. Chu, L. Maliskova) for their assistance with 10X Chromium and Visium library preparation; the Data science CoLab (especially E. Flynn) for their teaching and office hours in computational biology; the Parnassus Advanced Light Microscopy CoLab for their training and support in using the microscopes; and the Flow Cytometry CoLab for the guidance and preparation of cell sorters. We would also like to thank the Genomics group at CZ Biohub for their contributions to sequencing. Specific appreciation to A. Combes for his sharing of the Celleca cell counter used for counting cells and nuclei.

We would like to thank the members of the Malignant Germ Cell International Consortium (especially: T. Shioda, J. Amatruda) for their outstanding feedback to the project. We thank all members of the Laird lab (especially: E. Gaylord, J. Zussman, M. Foecke, T. Mcintyre, C. Doherty), all members of the MacKenzie lab (especially: B. Borges, T. Lum, K. Wedderburn-Pugh, A. Herzog), and our scientific colleagues (R. Samuel, J. Sandoval) for their thoughtful feedback on the manuscript. Thank you to P. Derish, director of the UCSF Surgery Scientific Writing Core for her editing and feedback. We would like to thank B. Panning and K. Shannon for guidance and help throughout this work on E.R. ’s thesis committee. And a thank you to S. Wamaitha, A. Clark, M. Sirota and S. Villeda for their mentorship to E.R. ’s PhD training. Schematics were created in BioRender (Publication License D. Laird (2025;))

## Author Contributions

E.J.R., T.C.M., D.J.L. conceived and designed the study. E.J.R., S.J.C., M.C. D.P. processed the samples for analyses. E.J.R., K.G., B.J.H. conducted the bioinformatics analyses and statistical analyses. S.J.C.,D.P., L.V., A.B., S.C.D. provided samples, clinical data, and helped with data curation as well as interpretation of results. E.J.R., K.G.,T.C.M., D.J.L. drafted the manuscript. All authors read and approved the manuscript prior to submission.

## Inclusion and ethics

We support inclusive, diverse, and equitable conduct of research. One or more authors of this paper received funding designed to increase minority representation in science. The UCSF and Colorado Children’s Hospital IRBs approved the snRNA-seq collections (UCSF IRB#: 10-00093; Colorado Children’s Hospital IRB#: 22-1906). Informed consent was waived for the spatial transcriptomic samples based on the IRB’s criteria that the research posed minimal risk, did not adversely affect subjects’ rights and welfare, was impracticable without the waiver, and additional pertinent information would be provided when appropriate (UCSF IRB#: 16-19356).

## Supplementary Materials

The accompanying Supplementary Material file contains Supplementary Figs. 1 to 5. Supplementary Tables 1 to 4 (Excel) are provided separately.

## References

1. Oosterhuis, J. W. & Looijenga, L. H. J. Human germ cell tumours from a developmental perspective. Nat. Rev. Cancer 19, 522–537 (2019).

2. Peiró, J. L., Sbragia, L., Scorletti, F., Lim, F. Y. & Shaaban, A. Management of fetal teratomas. Pediatr. Surg. Int. 32, 635–647 (2016).

3. Altman, R. P., Randolph, J. G. & Lilly, J. R. Sacrococcygeal teratoma: American Academy of Pediatrics Surgical Section Survey-1973. J. Pediatr. Surg. 9, 389–398 (1974).

4. Westerburg, B. et al. Sonographic prognostic factors in fetuses with sacrococcygeal teratoma. J. Pediatr. Surg. 35, 322–5; discussion 325 (2000).

5. Yoneda, A. et al. Impact of the histological type on the prognosis of patients with prenatally diagnosed sacrococcygeal teratomas: the results of a nationwide Japanese survey. Pediatr. Surg. Int. 29, 1119–1125 (2013).

6. van Heurn, L. J. et al. Prognostic accuracy of factors associated with poor outcome in prenatally diagnosed sacrococcygeal teratoma: A systematic review and meta-analysis. Prenat. Diagn. 43, 1495–1505 (2023).

7. Graf, J. L., Housely, H. T., Albanese, C. T., Adzick, N. S. & Harrison, M. R. A surprising histological evolution of preterm sacrococcygeal teratoma. J. Pediatr. Surg. 33, 177–179 (1998).

8. Padilla, B. E. et al. Sacrococcygeal teratoma: late recurrence warrants long-term surveillance. Pediatr. Surg. Int. 33, 1189–1194 (2017).

9. Sepulveda-Rincon, L. P. et al. Determining the potency of primordial germ cells by injection into early mouse embryos. Dev. Cell 59, 695–704.e5 (2024).

10. Khan, S. A. & Theunissen, T. W. Modeling X-chromosome inactivation and reactivation during human development. Curr. Opin. Genet. Dev. 82, 102096 (2023).

11. Runyan, C., Gu, Y., Shoemaker, A., Looijenga, L. & Wylie, C. The distribution and behavior of extragonadal primordial germ cells in Bax mutant mice suggest a novel origin for sacrococcygeal germ cell tumors. Int. J. Dev. Biol. 52, 333–344 (2008).

12. Busch, C., Oppitz, M., Wehrmann, M., Schweizer, P. & Drews, U. Immunohistochemical localization of nanog and Oct4 in stem cell compartments of human sacrococcygeal teratomas. Histopathology 52, 717–730 (2008).

13. Busch, C. et al. Isolation of three stem cell lines from human sacrococcygeal teratomas. J. Pathol. 217, 589–596 (2009).

14. Hambraeus, M. et al. Differential activation of immune effector processes in mature compared to immature sacrococcygeal teratomas. Fetal Pediatr. Pathol. 41, 413–425 (2022).

15. Wu, H., Kirita, Y., Donnelly, E. L. & Humphreys, B. D. Advantages of Single-Nucleus over Single-Cell RNA Sequencing of Adult Kidney: Rare Cell Types and Novel Cell States Revealed in Fibrosis. J. Am. Soc. Nephrol. 30, 23–32 (2019).

16. Slyper, M. et al. A single-cell and single-nucleus RNA-Seq toolbox for fresh and frozen human tumors. Nat. Med. 26, 792–802 (2020).

17. Quan, F. et al. Annotation of cell types (ACT): a convenient web server for cell type annotation. Genome Med. 15, 91 (2023).

18. Cao, M. et al. Characterization of immature ovarian teratomas through single-cell transcriptome. Front. Immunol. 14, 1131814 (2023).

19. McDonald, D. et al. Defining the Teratoma as a Model for Multi-lineage Human Development. Cell 183, 1402–1419.e18 (2020).

20. Cao, J. et al. A human cell atlas of fetal gene expression. Science 370, (2020).

21. Hou, W. & Ji, Z. Assessing GPT-4 for cell type annotation in single-cell RNA-seq analysis. Nat. Methods 21, 1462–1465 (2024).

22. Pollen, A. A. et al. Molecular identity of human outer radial glia during cortical development. Cell 163, 55–67 (2015).

23. Lee, L. Mechanisms of mammalian ciliary motility: Insights from primary ciliary dyskinesia genetics. Gene 473, 57–66 (2011).

24. Dredge, B. K. & Jensen, K. B. NeuN/Rbfox3 nuclear and cytoplasmic isoforms differentially regulate alternative splicing and nonsense-mediated decay of Rbfox2. PLoS ONE 6, e21585 (2011).

25. Küspert, M., Hammer, A., Bösl, M. R. & Wegner, M. Olig2 regulates Sox10 expression in oligodendrocyte precursors through an evolutionary conserved distal enhancer. Nucleic Acids Res. 39, 1280–1293 (2011).

26. Keller, L., Werner, S. & Pantel, K. Biology and clinical relevance of EpCAM. Cell Stress 3, 165–180 (2019).

27. Rezzani, R., Franco, C., Franceschetti, L., Gianò, M. & Favero, G. A Focus on Enterochromaffin Cells among the Enteroendocrine Cells: Localization, Morphology, and Role. Int. J. Mol. Sci. 23, (2022).

28. Hirsch, D. et al. LGR5 positivity defines stem-like cells in colorectal cancer. Carcinogenesis 35, 849–858 (2014).

29. Kumar, A. et al. Specification and Diversification of Pericytes and Smooth Muscle Cells from Mesenchymoangioblasts. Cell Rep. 19, 1902–1916 (2017).

30. Paulsson, M. & Heinegård, D. Purification and structural characterization of a cartilage matrix protein. Biochem. J. 197, 367–375 (1981).

31. Costa, M. L., Escaleira, R., Cataldo, A., Oliveira, F. & Mermelstein, C. S. Desmin: molecular interactions and putative functions of the muscle intermediate filament protein. Braz. J. Med. Biol. Res. 37, 1819–1830 (2004).

32. Zammit, P. S. et al. Pax7 and myogenic progression in skeletal muscle satellite cells. J. Cell Sci. 119, 1824–1832 (2006).

33. Barnett, S. N. et al. An organotypic atlas of human vascular cells. Nat. Med. 30, 3468–3481 (2024).

34. Tsai, M., Valent, P. & Galli, S. J. KIT as a master regulator of the mast cell lineage. J. Allergy Clin. Immunol. 149, 1845–1854 (2022).

35. Szabo, P. A. et al. Single-cell transcriptomics of human T cells reveals tissue and activation signatures in health and disease. Nat. Commun. 10, 4706 (2019).

36. Rebuffet, L. et al. High-dimensional single-cell analysis of human natural killer cell heterogeneity. Nat. Immunol. 25, 1474–1488 (2024).

37. Chistiakov, D. A., Killingsworth, M. C., Myasoedova, V. A., Orekhov, A. N. & Bobryshev, Y. V. CD68/macrosialin: not just a histochemical marker. Lab. Invest. 97, 4–13 (2017).

38. Fenton, T. M. et al. Immune Profiling of Human Gut-Associated Lymphoid Tissue Identifies a Role for Isolated Lymphoid Follicles in Priming of Region-Specific Immunity. Immunity 52, 557–570.e6 (2020).

39. De Leon-Oliva, D. et al. AIF1: Function and Connection with Inflammatory Diseases. Biology (Basel) 12, (2023).

40. Valdiserri, R. O. & Yunis, E. J. Sacrococcygeal teratomas: a review of 68 cases. Cancer 48, 217–221 (1981).

41. Xu, S. et al. Using clusterProfiler to characterize multiomics data. Nat. Protoc. 19, 3292–3320 (2024).

42. Ma, Y. & Zhou, X. Spatially informed cell-type deconvolution for spatial transcriptomics. Nat. Biotechnol. 40, 1349–1359 (2022).

43. Nguyen, Q. H. et al. Single-cell RNA-seq of human induced pluripotent stem cells reveals cellular heterogeneity and cell state transitions between subpopulations. Genome Res. 28, 1053–1066 (2018).

44. Wen, L. & Tang, F. Human Germline Cell Development: from the Perspective of Single-Cell Sequencing. Mol. Cell 76, 320–328 (2019).

45. Irie, N. et al. SOX17 is a critical specifier of human primordial germ cell fate. Cell 160, 253–268 (2015).

46. Loda, A. & Heard, E. Xist RNA in action: Past, present, and future. PLoS Genet. 15, e1008333 (2019).

47. Kawakami, T. et al. Characterization of loss-of-inactive X in Klinefelter syndrome and female-derived cancer cells. Oncogene 23, 6163–6169 (2004).

48. Chen, X. & Zhang, J. The X to autosome expression ratio in haploid and diploid human embryonic stem cells. Mol. Biol. Evol. 33, 3104–3107 (2016).

49. Chitiashvili, T. et al. Female human primordial germ cells display X-chromosome dosage compensation despite the absence of X-inactivation. Nat. Cell Biol. 22, 1436–1446 (2020).

50. Tukiainen, T. et al. Landscape of X chromosome inactivation across human tissues. Nature 550, 244–248 (2017).

51. Patel, A. P. et al. Single-cell RNA-seq highlights intratumoral heterogeneity in primary glioblastoma. Science 344, 1396–1401 (2014).

52. Petropoulos, S. et al. Single-Cell RNA-Seq Reveals Lineage and X Chromosome Dynamics in Human Preimplantation Embryos. Cell 165, 1012–1026 (2016).

53. Sirchia, S. M. et al. Loss of the inactive X chromosome and replication of the active X in BRCA1-defective and wild-type breast cancer cells. Cancer Res. 65, 2139–2146 (2005).

54. Hiester, A., Nettersheim, D., Nini, A., Lusch, A. & Albers, P. Management, treatment, and molecular background of the growing teratoma syndrome. Urol. Clin. North Am. 46, 419–427 (2019).

55. Norris, H. J., Zirkin, H. J. & Benson, W. L. Immature (malignant) teratoma of the ovary.A clinical and pathologic study of 58 cases. *Cancer* **37**, 2359–2372 (1976).

56. Roidor, C. et al. Temporal and regional X-linked gene reactivation in the mouse germline reveals site-specific retention of epigenetic silencing. Nat. Struct. Mol. Biol. 32, 926–939 (2025).

57. Sinnock, K. L., Perez-Atayde, A. R., Boynton, K. A. & Mutter, G. L. Clonal analysis of sacrococcygeal “teratomas”. Pediatr. Pathol. Lab. Med. 16, 865–875 (1996).

58. Li, J. et al. Long noncoding RNA XIST: Mechanisms for X chromosome inactivation, roles in sex-biased diseases, and therapeutic opportunities. Genes Dis. 9, 1478–1492 (2022).

59. Cardenas-Diaz, F. L. et al. Temporal and spatial staging of lung alveolar regeneration is determined by the grainyhead transcription factor Tfcp2l1. Cell Rep. 42, 112451 (2023).

60. Ye, T. et al. Nr5a2 promotes cancer stem cell properties and tumorigenesis in nonsmall cell lung cancer by regulating Nanog. Cancer Med. 8, 1232–1245 (2019).

61. Chen, W. & Wang, Y.-J. Multifaceted roles of OCT4 in tumor microenvironment: biology and therapeutic implications. Oncogene (2025) doi:10.1038/s41388-025-03408-x.

62. Janesick, A. et al. High resolution mapping of the tumor microenvironment using integrated single-cell, spatial and in situ analysis. Nat. Commun. 14, 8353 (2023).

63. Young, M. D. & Behjati, S. SoupX removes ambient RNA contamination from droplet-based single-cell RNA sequencing data. Gigascience 9, giaa151. (2020).

64. Hao, Y. et al. Dictionary learning for integrative, multimodal and scalable single-cell analysis. Nat. Biotechnol. 42, 293–304 (2024).

65. Marsh, S. E. scCustomize: Custom Visualizations & Functions for Streamlined Analyses of Single Cell Sequencing. Zenodo (2024) doi:10.5281/zenodo.14529706.

66. McGinnis, C. S., Murrow, L. M. & Gartner, Z. J. DoubletFinder: Doublet Detection in Single-Cell RNA Sequencing Data Using Artificial Nearest Neighbors. Cell Syst. 8, 329–337.e4 (2019).

67. Germain, P.-L., Lun, A., Garcia Meixide, C., Macnair, W. & Robinson, M. D. Doublet identification in single-cell sequencing data using scDblFinder. F1000Res. 10, 979 (2021).

68. Amezquita, R. A. et al. Orchestrating single-cell analysis with Bioconductor. Nat. Methods 17, 137–145 (2020).

69. Zappia, L. & Oshlack, A. Clustering trees: a visualization for evaluating clusterings at multiple resolutions. Gigascience 7, giy083 (2018).

70. Miller, S. A. et al. LSD1 and Aberrant DNA Methylation Mediate Persistence of Enteroendocrine Progenitors That Support BRAF-Mutant Colorectal Cancer. Cancer Res. 81, 3791–3805 (2021).

71. Phipson, B. & Smyth, G. K. Permutation P-values should never be zero: calculating exact P-values when permutations are randomly drawn. Stat. Appl. Genet. Mol. Biol. 9, Article39 (2010).

72. Barkas, N. et al. Joint analysis of heterogeneous single-cell RNA-seq dataset collections. Nat. Methods 16, 695–698 (2019).

73. Larsson, L., Franzén, L., Ståhl, P. L. & Lundeberg, J. Semla: a versatile toolkit for spatially resolved transcriptomics analysis and visualization. Bioinformatics 39, (2023).

74. Durinck, S., Spellman, P. T., Birney, E. & Huber, W. Mapping identifiers for the integration of genomic datasets with the R/Bioconductor package biomaRt. Nat. Protoc. 4, 1184–1191 (2009).

75. Tomofuji, Y. et al. Quantification of escape from X chromosome inactivation with single-cell omics data reveals heterogeneity across cell types and tissues. Cell Genomics 4, 100625 (2024).

76. Huang, X. & Huang, Y. Cellsnp-lite: an efficient tool for genotyping single cells. Bioinformatics 37, 4569–4571 (2021).

77. Yang, H. & Wang, K. Genomic variant annotation and prioritization with ANNOVAR and wANNOVAR. Nat. Protoc. 10, 1556–1566 (2015).

78. Andreatta, M. & Carmona, S. J. UCell: Robust and scalable single-cell gene signature scoring. Comput. Struct. Biotechnol. J. 19, 3796–3798 (2021).

79. Wang, R. H. & Thakar, J. Comparative analysis of single-cell pathway scoring methods and a novel approach. NAR Genom. Bioinform. 6, lqae124 (2024).

80. Subramanian, A. et al. Gene set enrichment analysis: a knowledge-based approach for interpreting genome-wide expression profiles. Proc Natl Acad Sci USA 102, 15545–15550 (2005).

81. Yu, G. et al. GOSemSim: an R package for measuring semantic similarity among GO terms and gene products. Bioinformatics 26, 976–978 (2010).

