## Supplement for "Integrated single-nuclei and spatial transcriptomic profiling of human sacrococcygeal teratomas reveals heterogeneity in cellular composition and X-chromosome inactivation"

**This PDF includes:**

Supplementary Figs. 1 to 5

**Other Supplementary Materials for this manuscript include the following:**

Supplementary Tables 1 to 4 (Excel) provided separately.

### Supplementary Figures

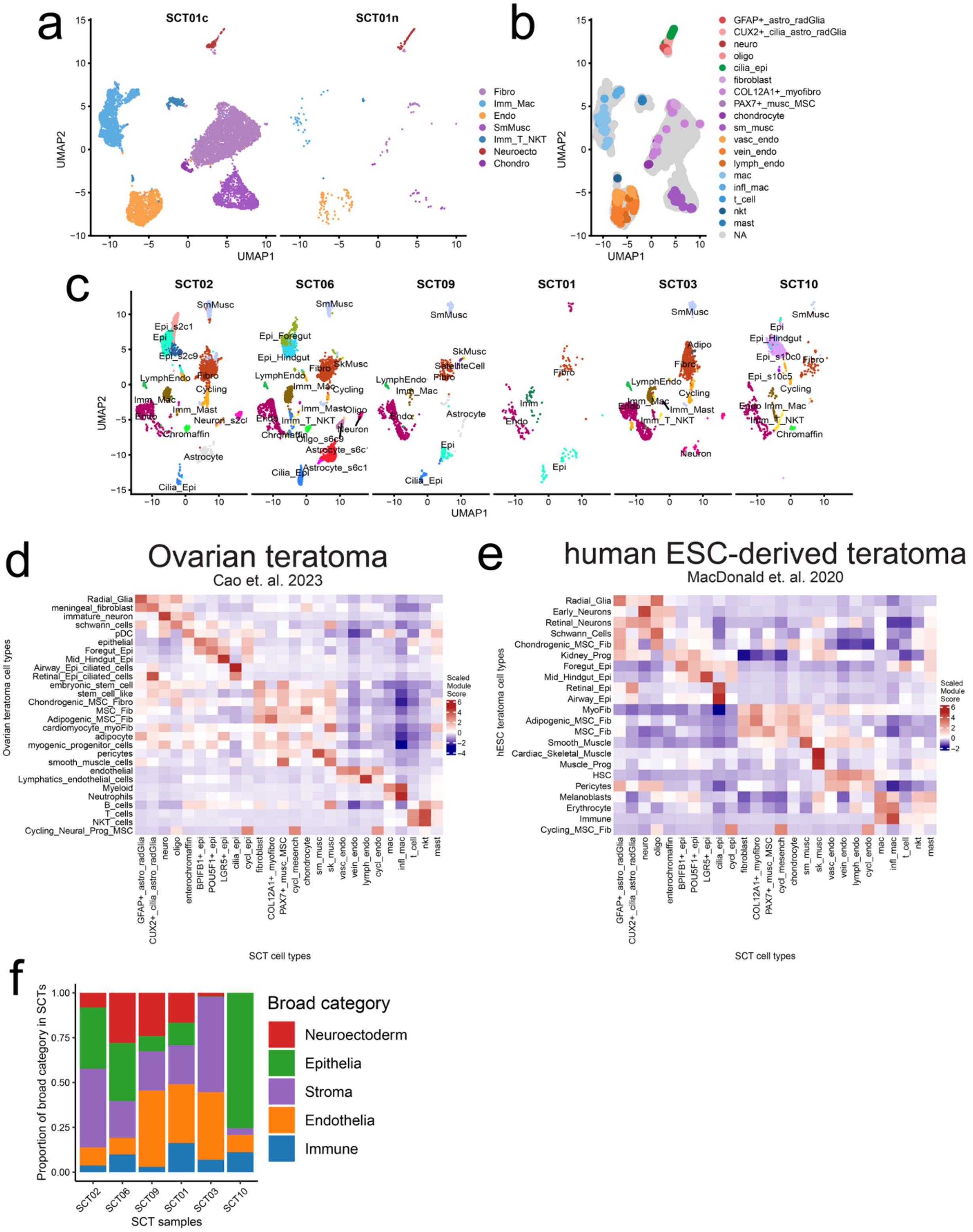


**Supplementary Fig. 1. Prior teratoma and developmental datasets informed annotation of cell types.**

(a) UMAP projection of integrated single-cell (SCT01c) and single-nuclei (SCT01n) samples taken from SCT01. (b) Overlap/projection of SCT01n on SCT01c. For technical reasons, SCT01n was left in -80C rather than liquid nitrogen, leading to low cell capture. However, of the cells captured there was broad overlap in cell identities between cells and nuclei in SCT01. We proceeded with nuclei for further analyses given nuclei captured all cell types independent of dissociation techniques. (c) Independent SCT annotations from each sample allowed for the naming of cell types among the individual tumors. We then mapped the cells from each cluster onto the same integrated UMAP space to validate overlapping identities. (d,e) Heatmap displaying the correlation between SCT sub-category clusters (x-axis) and ovarian teratoma clusters (y-axis) in d and human ESC-derived teratoma clusters (y-axis) in e. Red denotes high correlation and blue denotes low correlation. Most cell identities had considerable overlap, but there were some discrepancies in the names provided for the stromal population. (f) Bar plot colored by broad cell category from Figure 1b and grouped by SCT sample. SCT10 is the only sample missing a broad category - neuroectodermal lineage.

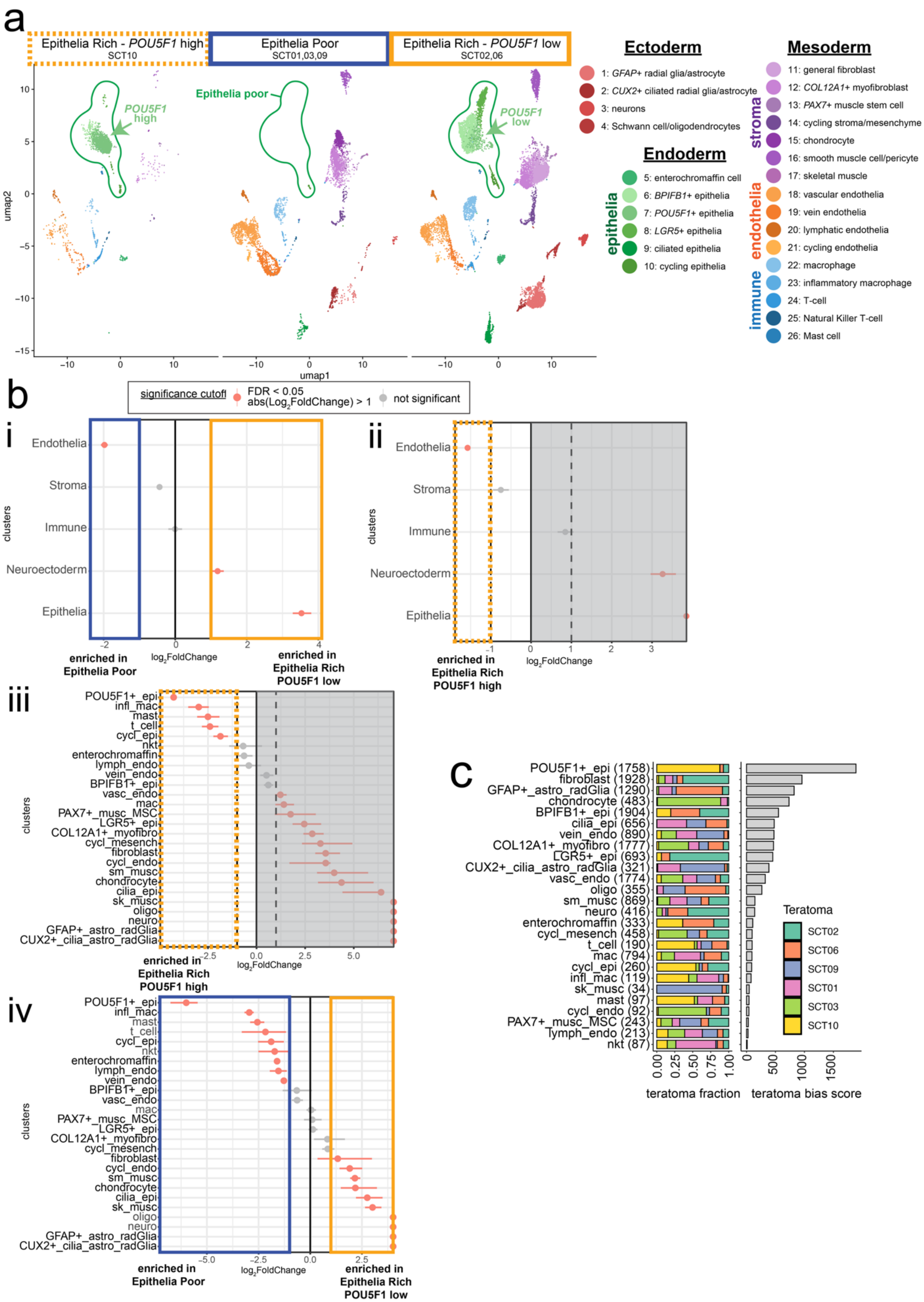
a
Epithelia Rich - POU5F1 high
SCT10
Epithelia Poor
SCT01,03,09
Epithelia Rich - POU5F1 low
SCT02,06
POU5F1 high
Epithelia poor
POU5F1 low
umap1
umap2
Ectoderm
1: GFAP+ radial glia/astrocyte
2: CUX2+ ciliated radial glia/astrocyte
3: neurons
4: Schwann cell/oligodendrocytes
Endoderm
epithelia
5: enterochromaffin cell
6: BPIFB1+ epithelia
7: POU5F1+ epithelia
8: LGR5+ epithelia
9: ciliated epithelia
10: cycling epithelia
Mesoderm
stroma
endothelia
immune
11: general fibroblast
12: COL12A1+ myofibroblast
13: PAX7+ muscle stem cell
14: cycling stroma/mesenchyme
15: chondrocyte
16: smooth muscle cell/pericyte
17: skeletal muscle
18: vascular endothelia
19: vein endothelia
20: lymphatic endothelia
21: cycling endothelia
22: macrophage
23: inflammatory macrophage
24: T-cell
25: Natural Killer T-cell
26: Mast cell
b
significance cutoff
FDR < 0.05
abs(Log2FoldChange) > 1
not significant
i
Endothelia
Stroma
Immune
Neuroectoderm
Epithelia
clusters
log2FoldChange
enriched in Epithelia Poor
enriched in Epithelia Rich POU5F1 low
ii
enriched in Epithelia Rich POU5F1 high
iii
POU5F1+_epi
infl_mac
mast
t_cell
cycl_epi
nkt
enterochromaffin
lymph_endo
vein_endo
BPIFB1+_epi
vasc_endo
mac
PAX7+_musc_MSC
LGR5+_epi
COL12A1+_myofibro
cycl_mesench
fibroblast
cycl_endo
sm_musc
chondrocyte
cilia_epi
sk_musc
oligo
neuro
GFAP+_astro_radGlia
CUX2+_cilia_astro_radGlia
enriched in Epithelia Rich POU5F1 high
iv
enriched in Epithelia Poor
enriched in Epithelia Rich POU5F1 low
c
POU5F1+_epi (1758)
fibroblast (1928)
GFAP+_astro_radGlia (1290)
chondrocyte (483)
BPIFB1+_epi (1904)
cilia_epi (656)
vein_endo (890)
COL12A1+_myofibro (1777)
LGR5+_epi (693)
CUX2+_cilia_astro_radGlia (321)
vasc_endo (1774)
oligo (355)
sm_musc (869)
neuro (416)
enterochromaffin (333)
cycl_mesench (458)
t_cell (190)
mac (794)
cycl_epi (260)
infl_mac (119)
sk_musc (34)
mast (97)
cycl_endo (92)
PAX7+_musc_MSC (243)
lymph_endo (213)
nkt (87)
Teratoma
SCT02
SCT06
SCT09
SCT01
SCT03
SCT10
teratoma fraction
teratoma bias score

**Supplementary Fig. 2. Characterizing the proportional differences in cell types among the three SCT clusters.**

(a) UMAP projection of the 3 categories of SCTs, (1) SCT10 - epithelia-rich *POU5F1*$^{high}$, (2) SCT01, 03, 09 - epithelia-poor, and (3) SCT02,06 - epithelia-rich *POU5F1*$^{low}$. Green arrow pointing to the *POU5F1+* population that is enriched in SCT10 (epithelia-rich POU5F1high) and is present in lower proportion in epithelia-rich *POU5F1*$^{low}$. Green circle shows the missing epithelia population in epithelia-poor samples. (b) Proportion testing for cell type enrichment across the 3 principal component (PC) clusters. Orange dots are significantly enriched with false discovery rate < 0.05 and absolute $log_2$Fold Change > 1 by Monte-Carlo/permutation test. Boxes around the enriched results correspond to their PC cluster: dashed-orange box is epithelia-rich *POU5F1*$^{high}$; solid-blue box is epithelia-poor; solid-orange box is epithelia-rich *POU5F1*$^{low}$; greyed out area is the comparison for epithelia-rich *POU5F1*$^{high}$. (i) We show broad cell types for epithelia-rich *POU5F1*$^{high}$ as compared to the rest of the samples and (ii) epithelia-poor as compared to epithelia-rich *POU5F1*$^{low}$ SCTs. (iii-iv) The same comparisons for the sub-category cell types enriched in each of the SCT samples. (c) On the left: the proportional makeup of each sample across cell type, with total cell number in parentheses. On the right: the bias score for each cell type shows bias towards specific teratomas. A low bias score means the cell type is well mixed across all 7 teratomas, whereas a high bias score means that the identity is made up by mostly one or few SCT samples. The highest bias score was in *POU5F1+* epithelia and it was made up of 87% from SCT10, 8% from SCT02, 5% from SCT06, 0% from SCT01, 03, and 09.

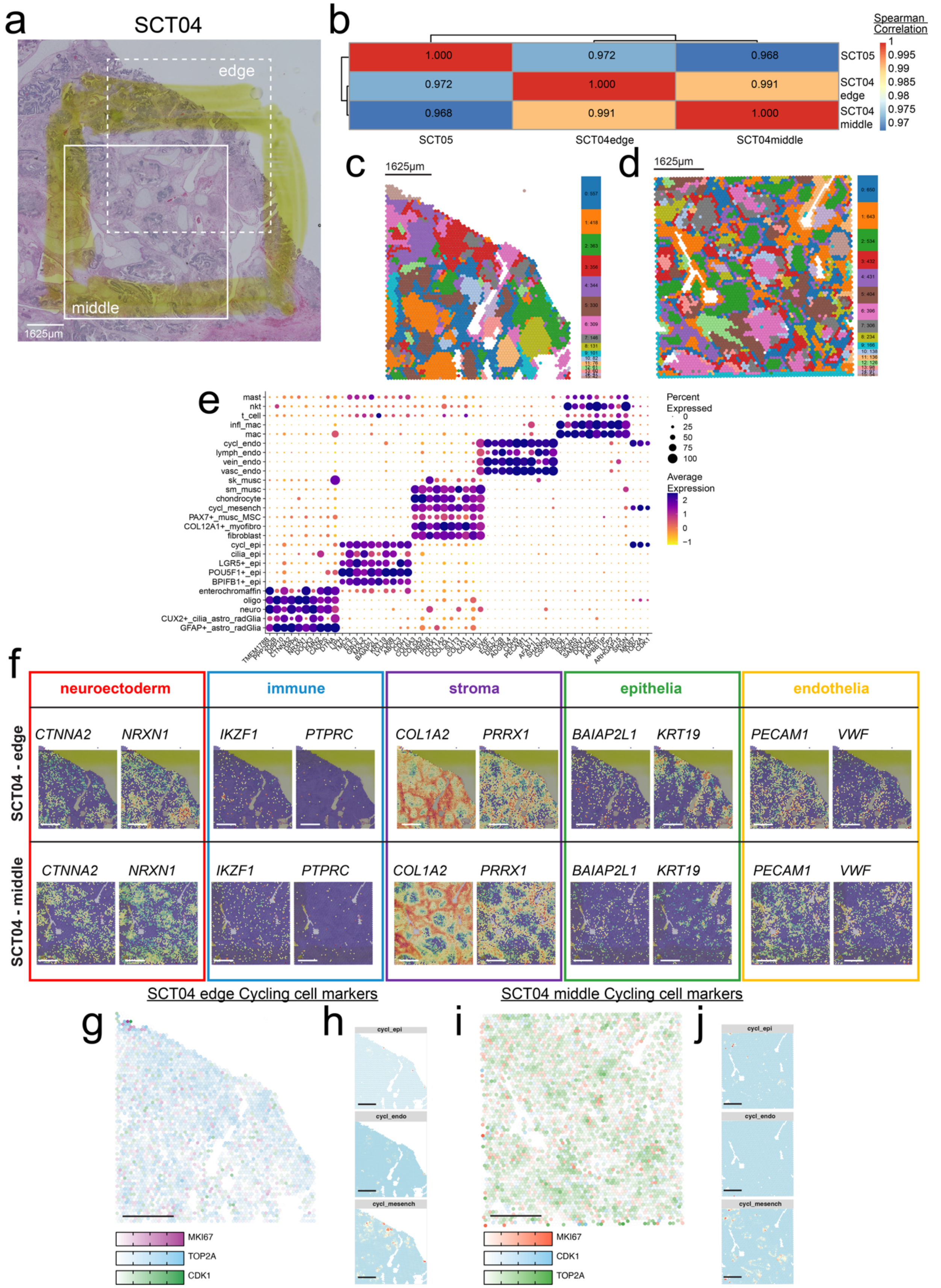


**Supplementary Fig. 3. Spatial transcriptomics of prenatal SCTs solid tumors.**

(a) Hematoxylin and eosin staining of prenatally-resected SCT (prSCT), SCT04, with box showing what areas denote the “edge” and “middle” spatial datasets. (b) Spearman correlation reveals that the two spatial datasets from SCT04 correlate more strongly with each other than with SCT05. Clustering was done by Euclidean distance. (c,d) Unbiased clusters overlaid in spatial plot for SCT04 edge (c) and middle (d). (e) Dot plot of key marker genes for each of the five broad categories of cell types in the snRNA-seq atlas. (f) Overlay of broad category markers taken from snRNA-seq atlas. Boxes correspond to the colors from Figure 1c, neuroectoderm in red, immune in blue, stroma in purple, epithelia in green, and endothelia in orange. (g,i) Cycling cell genes expressed across the SCT samples. Colors correspond to each gene: red = *MKI67*, green = *TOP2A*, blue = *CDK1*, with white being no expression. (h,j) Spatial distribution of cycling cell subtype spatial intensities by deconvolution analyses. All scale bars = 1625μm.

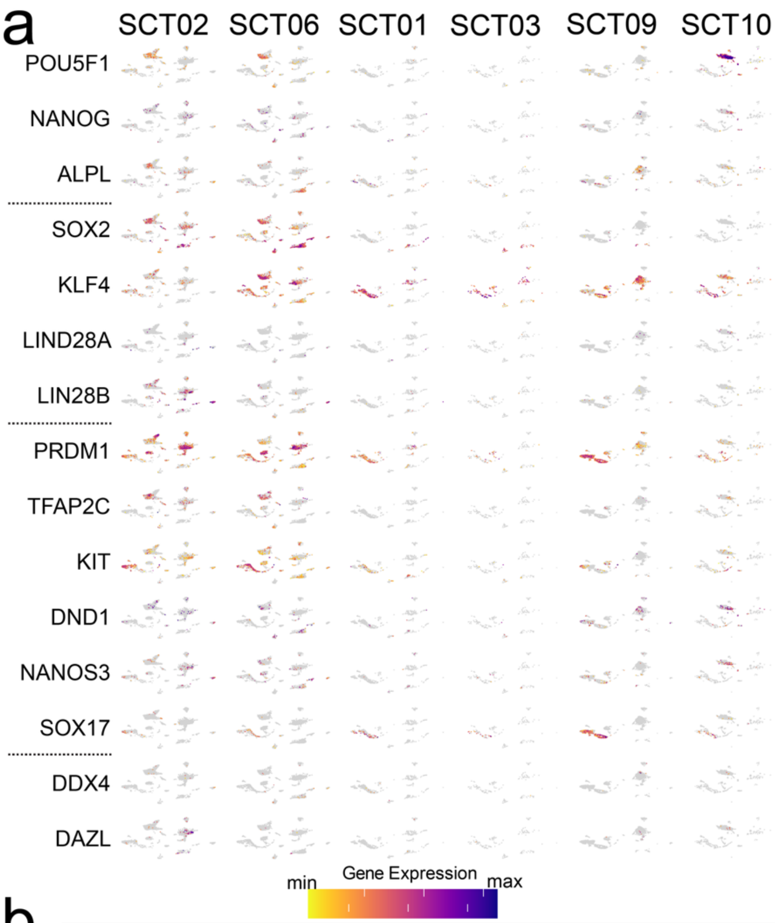


b

| *Genes expr. above threshold | SCT02 | SCT06 | SCT09 | SCT01 | SCT03 | SCT10 | Total cells |
|---|---|---|---|---|---|---|---|
| None | 5070 | 4579 | 803 | 199 | 2490 | 832 | 13973 |
| DAZL | 37 | 3 | 0 | 0 | 3 | 1 | 44 |
| DDX4 | 2 | 3 | 0 | 0 | 4 | 0 | 9 |
| NANOG | 8 | 15 | 3 | 0 | 2 | 1 | 29 |
| POU5F1 | 46 | 76 | 2 | 0 | 1 | 1152 | 1277 |
| PRDM1 | 734 | 454 | 69 | 14 | 384 | 53 | 1708 |
| SOX17 | 11 | 12 | 24 | 13 | 75 | 10 | 145 |
| SOX2 | 167 | 368 | 26 | 25 | 7 | 0 | 593 |
| TFAP2C | 47 | 51 | 0 | 0 | 5 | 0 | 103 |
| DAZL & PRDM1 | 6 | 0 | 0 | 0 | 0 | 0 | 6 |
| DAZL & SOX2 | 1 | 2 | 0 | 0 | 0 | 0 | 3 |
| DAZL & TFAP2C | 1 | 0 | 0 | 0 | 0 | 0 | 1 |
| DDX4 & POU5F1 | 0 | 0 | 0 | 0 | 0 | 1 | 1 |
| DDX4 & PRDM1 | 1 | 0 | 0 | 0 | 0 | 0 | 1 |
| DDX4 & SOX2 | 0 | 1 | 1 | 0 | 0 | 0 | 2 |
| NANOG & POU5F1 | 1 | 0 | 0 | 0 | 0 | 1 | 2 |
| NANOG & PRDM1 | 3 | 2 | 0 | 0 | 0 | 0 | 5 |
| NANOG & SOX2 | 0 | 0 | 1 | 0 | 0 | 0 | 1 |
| POU5F1 & PRDM1 | 2 | 0 | 1 | 0 | 0 | 5 | 8 |
| POU5F1 & SOX2 | 0 | 3 | 0 | 0 | 0 | 0 | 3 |
| POU5F1 & TFAP2C | 4 | 3 | 0 | 0 | 0 | 3 | 10 |
| PRDM1 & SOX17 | 1 | 2 | 8 | 2 | 59 | 5 | 77 |
| PRDM1 & SOX2 | 12 | 6 | 0 | 0 | 0 | 0 | 18 |
| PRDM1 & TFAP2C | 7 | 0 | 0 | 0 | 0 | 0 | 7 |
| SOX2 & TFAP2C | 3 | 3 | 0 | 0 | 0 | 0 | 6 |
| DAZL & POU5F1 & PRDM1 | 0 | 0 | 0 | 0 | 0 | 1 | 1 |
| PRDM1 & SOX2 & TFAP2C | 0 | 1 | 0 | 0 | 0 | 0 | 1 |

*Genes tested: POU5F1, NANOG, SOX2, SOX17, PRDM1, TFAP2C, DDX4, DAZL

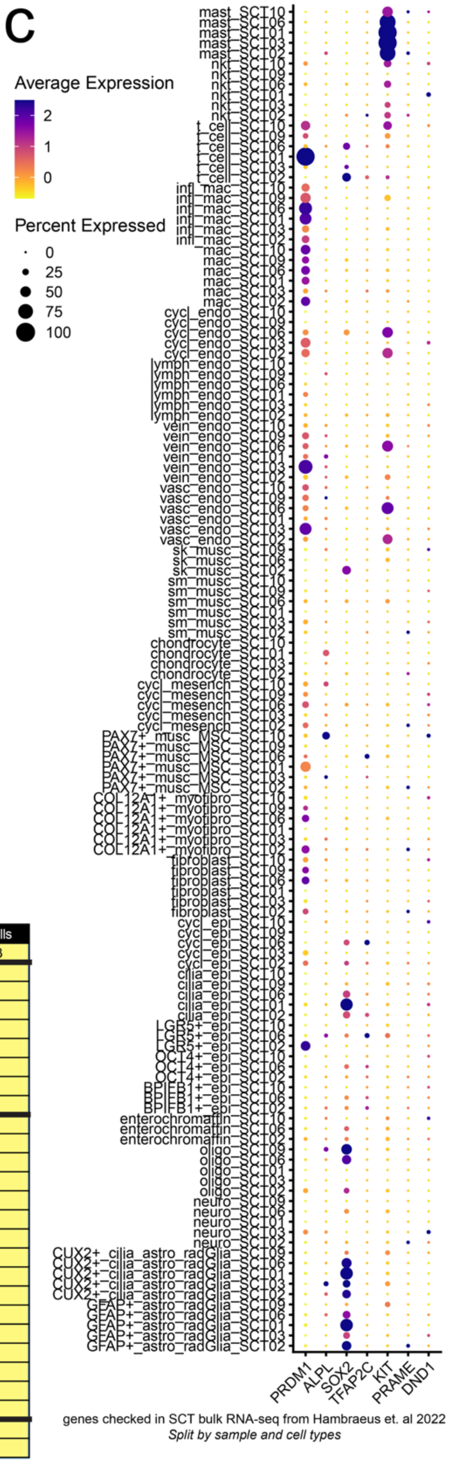


**Supplementary Fig. 4. Detailed mapping of genes found to be overexpressed in SCTs.**

(a) Feature plot showing the expression of each of the pluripotent and PGC genes from Figure 4a. (b) Combinatorial expression of key pluripotent and PGC master regulators: *POU5F1, NANOG, SOX2, SOX17, PRDM1, TFAP2C, DDX4, DAZL*. Dark lines distinguish cells that express no combinations, single-positive, double-positive, and triple-positive cells. On the right highlighted in yellow is the total cells that express each combination above the threshold of normalized expression > 0.5. (c) Dot plot of genes that were described as expressed in bulk RNA-seq of SCTs expanded from Figure 4B split by sub-category cell types. Scale bars = 1625μm.

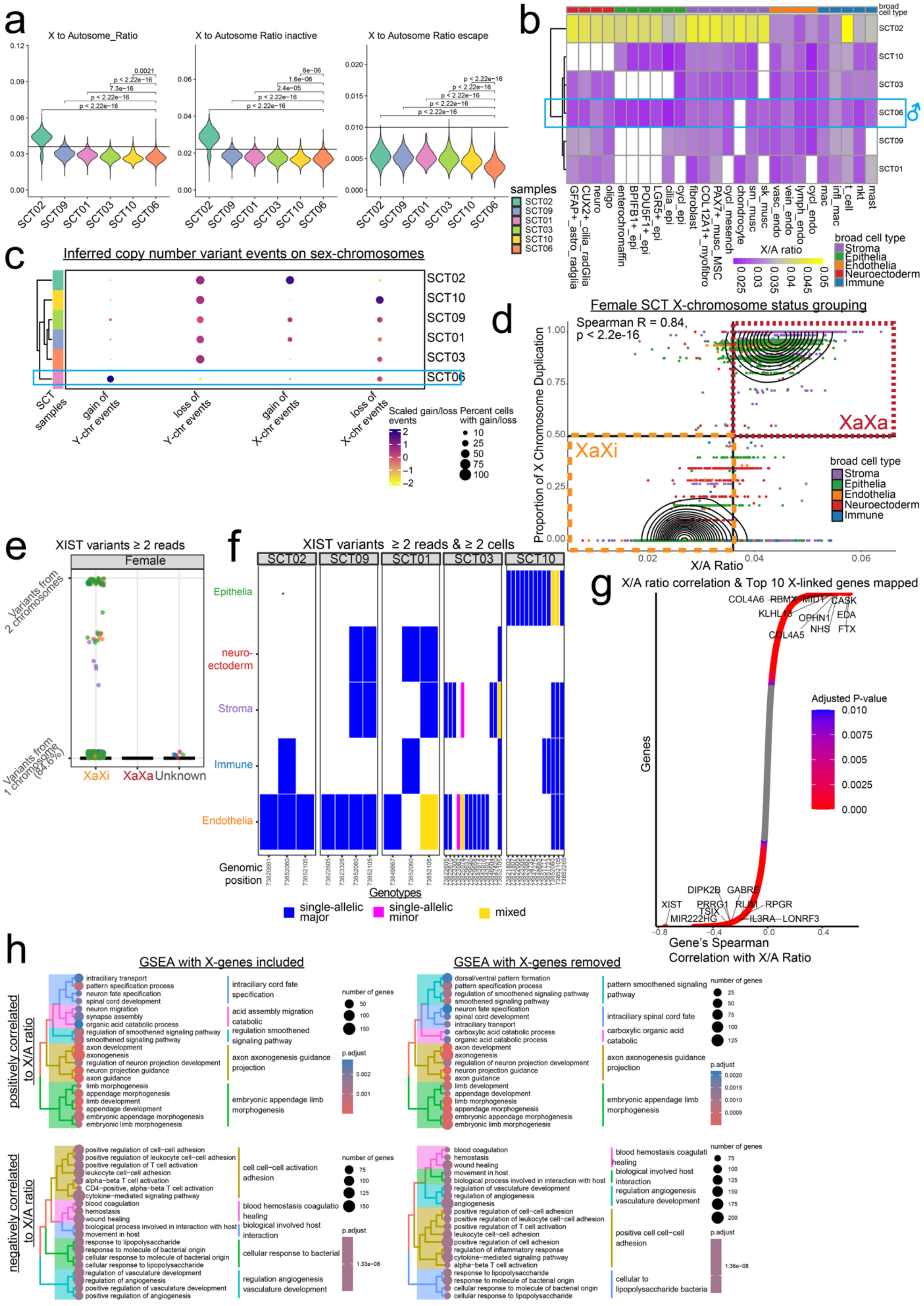
a
X to Autosome_Ratio
X to Autosome Ratio inactive
X to Autosome Ratio escape
SCT02 SCT09 SCT01 SCT03 SCT10 SCT06
samples
b
broad cell type
X/A ratio
broad cell type
Stroma
Epithelia
Endothelia
Neuroectoderm
Immune
c
Inferred copy number variant events on sex-chromosomes
SCT samples
gain of Y-chr events
loss of Y-chr events
gain of X-chr events
loss of X-chr events
Scaled gain/loss events
Percent cells with gain/loss
d
Female SCT X-chromosome status grouping
Spearman R = 0.84, p < 2.2e-16
Proportion of X Chromosome Duplication
X/A Ratio
XaXa
XaXi
e
XIST variants ≥ 2 reads
Female
Variants from 2 chromosomes
Variants from 1 chromosome (84.6%)
XaXi XaXa Unknown
f
XIST variants ≥ 2 reads & ≥ 2 cells
Epithelia
neuro-ectoderm
Stroma
Immune
Endothelia
Genomic position
Genotypes
single-allelic major
single-allelic minor
mixed
g
X/A ratio correlation & Top 10 X-linked genes mapped
Genes
Adjusted P-value
Gene's Spearman Correlation with X/A Ratio
h
GSEA with X-genes included
GSEA with X-genes removed
positively correlated to X/A ratio
negatively correlated to X/A ratio
number of genes
p.adjust

85

**Supplementary Fig. 5. Characterization of the X-chromosome gene expression for female SCTs.**

(a) X-to-Autosome (X/A) ratio violin plots stratified by SCT samples. The three plots are then separated by overall X/A ratio, X/A ratio for X-linked genes that are inactivated during X inactivation (XCI), and X/A ratio for X-linked genes that are escapees XCI. (b) Heatmap plot of the SCT samples (rows) and the sub-category cell types (columns). Purple shows low X/A ratio (< 0.036), yellow indicates high X/A ratio (> 0.036), with grey being 0.036. Broad cell types are highlighted at the top of the columns. (c) Dot plot showing inferred gain and loss events on the sex chromosomes from inferCNV with the male sample highlighted as successfully showing presence of Y-chromosome. (d) Scatter plot showing a significant positive correlation between the X/A ratio and the proportion of the X that is duplicated ($R=0.84$, $p<0.01$). We used a cutoff of X/A ratio > 0.036 and proportion of the X duplicated > 0.5 to call cells that we suspect have two active Xs (XaXa-like, burgundy box) versus those that have one active X (XaXi, orange box). Cells that did not land in these quadrants were designated as “unknown”. Density plots are overlaid to show where the cells are most concentrated. (e) Genotyping of single nucleotide polymorphisms (SNPs) on single-nuclei RNA molecules. We determined whether reads originated from one or both Xs in each cell. Dots represent SNP positions from *XIST* RNA split by XaXa-like, XaXi, and unknown cells on the x-axis. Dots are colored by broad category. Y-axis represents how much of each SNP came from one X or both Xs. In XaXi cells, 84.6% of *XIST* variants come from only one X. (f) Single X variants of *XIST* plotted in tiles by sample (broad grey boxes) and genomic position (x-axis). Single-allelic major (blue tiles) refers to the most abundant genotype at that position. Single-allelic minor (magenta tiles) refers to the alternative to single-allelic major. Mixed (yellow) refers to positions where some cells have alleles from both genotypes. (g) Genes plotted in ranked order by Spearman correlation to the X/A ratio. Highlighted are the top and bottom ten X-linked genes, highlighting that there are X-linked genes that strongly positively and negatively correlate with X/A ratio. Colors are the adjusted p-values ranging from 0 in red to 0.01 in blue; grey values are not considered significantly correlated. (h) Tree plot of semantically simplified gene set terms for top 20 gene ontology terms by adjusted p-value. Circle sizes are the number of genes in the term and colored by adjusted p-value.

### **Supplementary Tables** *(in Excel)*

**Supplementary Table 1. Orthogonal naming methods results for independent SCT samples**

(a) Naming results based on ACT automated cell annotation via a hierarchically organized marker gene map (b) Results of naming by correlated clusters with published ovarian (column "CellName_OvTe") and human embryonic stem cell (ESC)-derived teratoma (column "CellName_CellDerived") datasets (c) Names by using Seurat's RunAzimuth, running the "fetusref" from Cao et al. (2020) and retaining the top ("Percentage") average score ("AverageScore") for each cluster. (d) Naming using a large language model-based framework where the top 50 genes from each cluster were provided to ChatGPT's gpt-4o model.

**Supplementary Table 2. Marker genes for each cluster.**

Markers for all 26 cell types in SCTs using the Wilcoxon Rank Sum test via the FindAllMarkers function (logfc.threshold = 0.25, only.pos = TRUE).

**Supplementary Table 3. Principle component analysis genes and gene set results.**

(a) Gene loadings that define principle component 1 ranked by highest to lowest contributions. (b) Gene set enrichment analyses results using gene ontology for PC1 ranked genes by loading contribution. (c) Gene loadings that define principle component 2 ranked by highest to lowest contributions. (d) Gene set enrichment analyses results using gene ontology for PC2 ranked genes by loading contribution.

**Supplementary Table 4. Results of X-to-Autosomal ratio profiling using Spearman correlations and gene set enrichment.**

(a) Gene correlations to X-to-Autosomal Ratio using Spearman correlation and to account for multiple comparisons, p-values were adjusted using the Benjamini-Hochberg FDR correction (qvalue) (b) Gene set enrichment analyses results using gene ontology for all genes (autosomal and X-linked) (c) Gene set enrichment analyses results using gene ontology using only the autosomal genes.
